# Benchmarking long-read RNA-seq across modalities, methods, and sequencing depth in iNeurons

**DOI:** 10.64898/2026.04.01.715783

**Authors:** Gianfranco Botta, David Wissel, Samuel Higgins, Tomasz Chelmicki, Alexander Popescu, Kalina Radoynovska, Seraphin Probst, Julieta Ramírez Cuéllar, Kim Schneider, Zeynab-Mitra Nayernia, Ashley Byrne, Christopher D. Nelson, Zora Modrusan, Stormy J. Chamberlain, William Stephenson, Mark D. Robinson, Rajib Schubert

**Author notes:** Equal contribution (Gianfranco Botta, David Wissel). Equal supervision (William Stephenson, Mark D. Robinson, Rajib Schubert). Current affiliation (Rajib Schubert).

## Abstract

Long-read RNA sequencing (lrRNA-seq) provides advantages for transcript discovery and quantification through the sequencing of full-length transcripts. Although recent benchmarks have evaluated long-read technologies and quantification tools, to the best of our knowledge, no study to date has jointly compared sequencing technology, quantification choice, and depth across both bulk and single-cell platforms. Here, we generate a matched dataset using NGN2-induced neurons derived from Fragile X syndrome and isogenic rescue lines, profiled with bulk and single-cell Illumina, Oxford Nanopore Technologies (ONT), and Pacific Biosciences (PB) Kinnex technologies. All platforms and technologies capture the expected FMR1 reactivation signal. We find that PB bulk under-detects and under-quantifies short transcripts (less than 1.25 kb), ONT bulk under-detects and under-quantifies long transcripts (greater than 5 kb), and single-cell long-read technologies a large number of single-cell specific transcripts associated with truncations. Across six bulk and four single-cell long-read quantification tools, Isosceles, Miniquant, and Oarfish provide the best compromise between computational efficiency and performance in terms of quantification accuracy as measured by spike-ins, comparisons to Illumina, and on consensus based down-stream tasks such as differential transcript expression (DTE). Depth-equivalency analyses reveal that PB single-cell sequencing requires approximately three- to four-fold greater depth than bulk to reach comparable power for transcript discovery and differential transcript expression. Our work aims to offer practical guidance for study design, including the choice of technology, sequencing depth, and quantification method. In addition, we hope our data may serve a reference dataset to evaluate emerging long-read transcriptomic protocols and methods as well as more closely investigate FMR1 biology.

## Introduction

Technologies for long-read RNA-seq (lrRNA-seq) are transforming the study of gene regulation by enabling the capture of full-length transcripts (Monzó et al. 2025). Previous research has shown outstanding performance of lrRNA-seq compared to short-read RNA-seq (srRNA-seq) on transcript discovery and for the quantification of transcript-level expression (Wissel et al. 2026; Chen et al. 2025a; You et al. 2025; Pardo-Palacios et al. 2024; Chen et al. 2025b; Dong et al. 2023; Su et al. 2024).

Recently, lrRNA-seq has also been applied to single-cell experiments, demonstrating applications such as cell type-specific splicing, mutation detection, and immune repertoire profiling (Gupta et al. 2024). As a result, recent years have seen a surge in new computational methods to perform transcript quantification and discovery with lrRNA-seq (Chen et al. 2023; Prjibelski et al. 2023; Kabza et al. 2024; Loving et al. 2025; Zare Jousheghani et al. 2025; Li et al. 2025; Kovaka et al. 2019; Tang et al. 2024). While recent work has taken to benchmarking these methods, most prior work is focused on a subset of platforms or technologies (Wissel et al. 2026; Dong and Ritchie 2023; Chen et al. 2025a; Pardo-Palacios et al. 2024). Other comparisons encompass an exhaustive array of platforms but do not consider different options for computational methods to analyze them (Dong and Ritchie 2023; Chen et al. 2025a), or are focused on one particular task (Kabza et al. 2024). In what follows, we briefly survey relevant literature as far as it pertains to the benchmarking or comparison of lrRNA-seq technologies and/or methods. We note that we focus primarily on quantification and related downstream tasks such as differential transcript expression (DTE) and differential transcript usage (DTU).

Dong and Ritchie (2023) leveraged triplicates of lung adenocarcinoma cell lines to benchmark the performance of bulk Oxford Nanopore Technologies (ONT) cDNA lrRNA-seq and Illumina srRNA-seq, additionally leveraging in silico mixtures to create ground truths in addition to sequin spike-ins. The authors showed that while StringTie2 and Bambu performed well for transcript discovery on their ONT dataset, the choice for DTE and, especially DTU methods was less clear, with several methods performing equally well. Chen et al. (2025b) sequenced seven human cell lines, each with multiple replicates, with several bulk lrRNA-seq technologies, including Pacific Biosciences (PB) Iso-Seq, ONT cDNA (with PCR), ONT cDNA (without PCR), and ONT direct RNA (among others). The authors show that different technologies and/or protocols produce notable differences in terms of depth, read length, coverage, and other aspects. In addition, while the gene-level quantifications between the various lrRNA-seq technologies and srRNA-seq showed good concordance, they show that transcript-level analyses generally benefit from the improved read-to-transcript assignment possible with lrRNA-seq. The LRGASP project ran a community challenge with the goal of determining the performance of both technologies and methods for transcript discovery, transcript quantification, and de novo transcript discovery from bulk lrRNA-seq (Pardo-Palacios et al. 2024). The authors generated data for human, mouse, and manatee, prepared four library protocols (cDNA, direct RNA, R2C2, and CapTrap), and sequenced data using sequencers from PB, ONT, and Illumina. Method developers could submit to one or several challenges for one or all data types. The challenge and subsequent analysis revealed that Isoquant, FLAIR, and Bambu were among the top performers for transcript quantification on both ONT cDNA and PB. Wissel et al. (2026) compared high-depth bulk Kinnex PB lrRNA-seq of an iPSC to endothelial differentiation to comparably deep Illumina srRNA-seq and concluded that PB enables improved performance on quantification-related transcript-level tasks due to its improved performance in resolving read-to-transcript assignment ambiguity. Recently, Chen et al. (2025a) leveraged Quartet reference materials to sequence one of the largest existing bulk lrRNA-seq datasets, incorporating PB Kinnex, PB Iso-Seq, ONT cDNA, and ONT direct RNA, as well as srRNA-seq. They use their frame-work to construct a ground truth dataset which reveals that quantification methods combining lrRNA-seq and srRNA-seq perform especially well.

On the single-cell front, Tian et al. (2021) developed one of the first modified Chromium 10x protocols that enables joint sequencing of single-cell lrRNA-seq and srRNA-seq of the same cells. The authors also developed a method, FLAMES, that performs transcript discovery and other long-read specific tasks in single-cell RNA-seq (RNA-seq). Later, Zajac et al. (2025) ran a quality evaluation study comparing single-cell PB Kinnex RNA-seq to single-cell srRNA-seq. They show that while globally, single-cell Kinnex lrRNA-seq and single-cell srRNA-seq produced largely concordant results in terms of gene expression quantifications and detected cells, single-cell Kinnex lrRNA-seq provides much more control over different sequencing artefacts commonly encountered in 10X data. Recently, Scoones et al. (2025) presented a comparison of several single-cell lrRNA-seq technologies, including ONT cDNA, PB Kinnex (sequenced on both the Revio and the Sequel IIe), as well as Illumina single-cell RNA-seq. Furthermore, the authors included the CRISPRclean kit by Jumpcode Genomics in both PB samples sequenced on the Revio and the Sequel IIe (meaning, both sequencers were used to sequence both libraries with and without the CRISPRclean kit). Their work highlights that while all single-cell lrRNA-seq techniques are broadly concordant in terms of quantification data, but highlights that the typical 10X Chromium library preparation results in molecules that can be quite short and not necessarily full-length. Furthermore, the authors show that common single-cell lrRNA-seq workflows result in count depth up to ten times lower than the raw count depth. You et al. (2025) recently presented LongBench, a single-cell and bulk lrRNA-seq dataset and benchmark that sequenced eight cancer cell lines using bulk lrRNA-seq as well as single-cell and singlenuclei lrRNA-seq. For bulk, the authors sequenced Illumina, PB Kinnex, ONT cDNA and ONT direct RNA, while for the single-nuclei and single-cell data, 10X Chromium libraries were sequenced using single-cell PB Kinnex and single-cell ONT cDNA. Their work evaluated quality control, quantification accuracy, and variant calling across all library-types and technologies. Most notably, they show that while gene-level analyses tend to be extremely concordant between libraries and technologies, transcript-level analyses can differ considerably, and more work is needed to resolve these differences.

The purpose of our work is to provide a comprehensive and neutral assessment of lrRNA-seq platforms, technologies, and computational methods for transcriptomic analysis using a neurodevelopmental model system. By systematically evaluating ONT and PB platforms across both bulk and single-cell technologies, we aim to clarify biases between technologies and platforms, understand which methods perform well for lrRNA-seq quantification, and what sequencing depths are necessary to achieve approximately equivalent outcomes in downstream tasks between platforms and technologies.

We chose a human neuronal model system to benchmark lrRNA-seq since neurons express a high number of long transcripts as well as widespread and highly conserved alternative splicing patterns, both of which place stringent requirements on the quality and biases of lrRNA-seq platforms and technologies (Zylka et al. 2015; Raj and Blencowe 2015).

## Results

### All platforms produced high-quality data that recapitulate FMR1 rescue

We generated bulk and single-cell RNA-seq datasets from NGN2-induced neurons under two conditions: a Fragile X syndrome line (E3) characterized by silencing of the FMR1 gene and the associated loss of expression phenotype, and a CRISPR-edited isogenic rescue line (IsoB11) in which targeted correction of the FMR1 locus restores FMR1 expression (Santoro et al. 2012; Maussion et al. 2023). The two conditions provide a system with a known loss-of-expression state and a well-characterized restoration state, enabling rigorous evaluation of cross-platform and technology performance using Illumina, ONT, and PB sequencing. Spike-in controls (ERCC and SIRV) were included in all bulk samples to facilitate ground-truth assessments. In all sequencing platforms and technologies, samples were sequenced deeply: for bulk Illumina sequencing, we obtained an average of 80.2M (million) raw reads per sample, compared to 48.2M for ONT and 22.1M for PB; for the single-cell format, Illumina returned an average of 296.2M reads, while ONT and PB had 105.1 and 100.9M reads, respectively (Fig. 1B). While we targeted 0.5% abundance each for the ERCC and SIRV spike-ins, actual recovery varied (computed as genome alignment proportions): Illumina retained 0.12% ERCC and 0.22% SIRV; ONT retained 0.076% ERCC and 0.61% SIRV; and PB retained 0.73% ERCC and 0.80% SIRV. Human genome alignment rates also varied, with Illumina and PB showing higher fractions (97.4% and 98.4%, respectively) than ONT (87.2%).

**Fig. 1:**
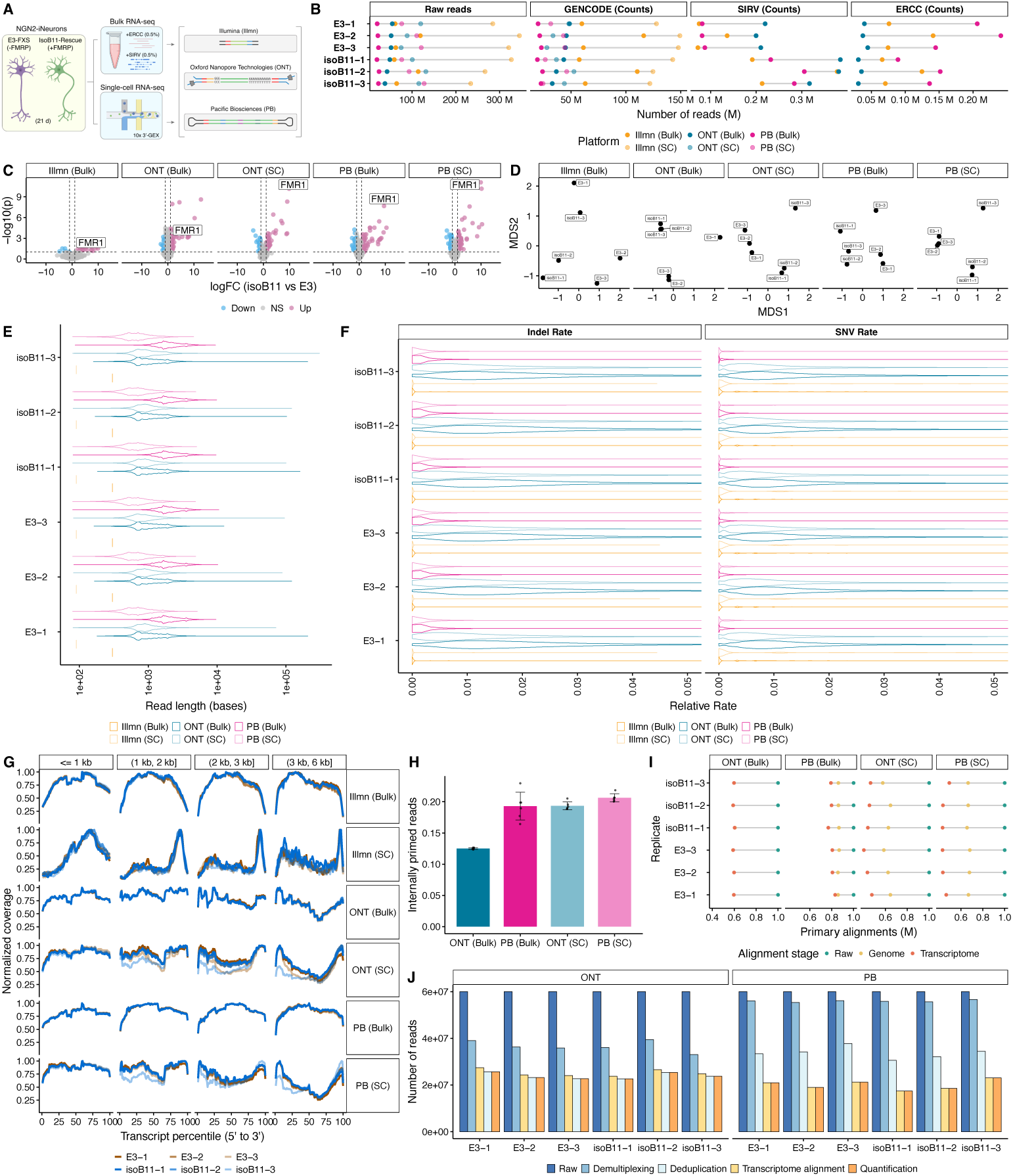
Quality control of the FXS cell lines system. **A.** Experimental design. NGN2-induced neurons derived from Fragile X syndrome (E3) and isogenic rescue (IsoB11), with FMRP reactivation, lines were profiled with Illumina, PB, and Oxford Nanopore technologies. For each line, three biological replicates were generated and sequenced using both bulk and single-cell technologies. SIRV and ERCC spike-ins were added to the bulk samples. **B**. Total number of raw reads and counts obtained for each sequencing platform. **C.** DGE analysis between E3 and IsoB11 samples. Bulk datasets were quantified with Isosceles, and pseudobulk single-cell datasets were quantified with Oarfish. In both analyses, FMR1 emerges as one of the top differentially expressed genes between conditions. **D.** MDS plots for each sequencing platform and technology based on transcript-level counts. Across all datasets, the primary axis of separation reflects the biological condition, with E3 and IsoB11 samples forming distinct clusters. **E.** Read length distributions for all datasets on 1M randomly sampled raw reads. Single-cell long-read sequencing platforms consistently produce shorter reads compared to their bulk counterparts. **F.** Relative error rates of insertions/deletions (Indel) and single-nucleotide variants (SNV) across sequencing platforms and technologies, obtained employing genome alignments (same reads as **E**). ONT data exhibits the highest error rates due to the intrinsic noisiness of the platform. **G.** Normalized (scaled 0–1) read coverage averaged across transcripts for each sequencing platform and technology (same reads as **E**). Single-cell datasets display more uneven coverage along transcripts compared to bulk datasets. **H.** Proportion of internally primed reads (see **Methods**) per technology and platform (same reads as **E**). Single-cell platforms consistently yield more internally primed reads than their bulk counterparts, although the difference for PB was only moderate. **I.** Portion of reads aligning to the human genome, and transcriptome, respectively (same reads as **E**). Long-read single-cell platforms consistently had less reads aligning to the genome and transcriptome than their bulk counterparts, pointing to lower efficiency. **J.** Progression of read counts through the analysis pipeline for single-cell long-read platforms. The total counts retained after pre-processing remain relatively low compared to the number of considered raw reads (60M). The order of deduplication and alignment steps differs between the two platforms due to discrepancies of the ONT and PB pre-processing pipelines.

Since we observed notable imbalances in read numbers both among samples within and across platforms, we downsampled all platforms and technologies to a common sequencing depth for further analysis. In particular, all bulk datasets were downsampled to 15M reads per sample, and all single-cell datasets to 60M reads per sample (see **Methods**). We also note that, hereafter, we primarily focus on *transcript-level* analyses, as far as quantification is concerned. Due to the inability of Illumina single-cell data to generate transcript-level counts, we exclude it from these analyses. Unless otherwise noted, all single-cell lrRNA-seq datasets were quantified using Oarfish, and all bulk lrRNA-seq datasets using Isosceles. Illumina data was quantified using Salmon and alevin-fry for bulk and single-cell technologies, respectively (see **Methods**).

Across all three platforms, both bulk and single-cell technologies confirmed the genetic rescue of FMR1 in the isogenic line, which was identified as one of the top differentially expressed genes, with all platforms showing broadly concordant gene-level log-fold changes (Fig. 1C, Additional file 1: Supplementary Fig. 1A). Further, FMR1 was consistently detected as one of the top most differentially expressed genes irrespective of the chosen quantification method (Additional file 1: Supplementary Fig. 1B-E). Overall, the quantifications of most platforms and technologies identified the main condition (IsoB11 vs E3) as the primary transcriptional difference between samples, as evidenced by the first multidimensional scaling (MDS) dimension, although Illumina bulk and ONT bulk were less clear than other platforms and technologies (Fig. 1D). MDS plots were congruent across quantification methods (Additional file 1: Supplementary Fig. 1F-I). Overall, within condition, all platforms and technologies were very replicable, with Pearson correlations consistently at 0.95 or above, although some quantification methods had slightly lower correlations around 0.92 (Additional file 1: Supplementary Fig. 1J-M). Between conditions, correlations were also consistently high, indicating limited transcriptional changes between E3 and isoB11 conditions, consistent with our analysis of differential gene expression (DGE) (Fig. 1C).

We first assessed read quality and length distributions across platforms. Both Illumina bulk and single-cell produced short fragments, as expected, with a mean length of 302 and 90 bp, respectively (Fig. 1E). Long-read platforms generated substantially longer reads, on average 2.1 kb and 1.2 kb for bulk PB and ONT, respectively. In general, long-read single-cell datasets generated shorter reads than their bulk counterparts, with an average length of 0.87 kb for single-cell PB and 1 kb for single-cell ONT.

Since PHRED values may not be calibrated across platforms and basecallers, we analyzed empirical error rates based on genome alignments. Overall, ONT bulk achieved an overall empirical error rate of 0.0345 (0.0140 SNV-related, 0.0210 for indels; Fig. 1F). Single-cell ONT behaved similarly, with an overall empirical error rate of 0.0264 (0.0156 SNVs and 0.0108 indels). PB bulk and Illumina bulk both had significantly lower overall empirical error rates, 0.00276 and 0.00315, respectively, driven primarily by their much lower indel-related error rates (0.00203 and 0.000161). The single-cell technology of PB and Illumina behaved similar to their bulk counterparts, achieving overall empirical error rates of 0.00276 and 0.00418, with indel- and SNV-related error rates at 0.00251, 0.000199, and 0.00185, 0.00398, respectively.

We then examined gene-level normalized coverage. With the exception of Illumina single-cell data, where methodological limitations restrict coverage to the 3’ end of each gene, we observed higher overall coverage in the bulk datasets (Fig. 1G). In contrast to bulk long-read data, long-read single-cell technologies displayed more prominent 3’ coverage for transcripts *>*2kb, potentially indicating suboptimal reverse transcription in the droplet format. Internal priming, as measured by PrimeSpotter (You et al. 2025) (see **Methods**), was moderate in all technologies and platforms, with ONT bulk exhibiting the lowest mean proportion of internally-primed reads (∼12.5%), while PB bulk was closer to the single-cell long-read technologies with ∼19.3%, and ONT single-cell gave ∼19.3% and PB single-cell ∼20.6% (Fig. 1H). To the best of our knowledge, PrimeSpotter’s effectiveness should not be impacted by the differing error profiles, given that its primary function is to mark reads as potentially internally priming if they either start with an unusually T-rich region or end with an unusually A-rich one.

In terms of on-target efficiency, measured by proportion of reads aligning to the genome, and transcriptome, respectively, PB bulk achieved the best performance. In particular, PB bulk had an average of ∼86.2% and ∼80.1% of reads aligning to the genome and transcriptome, respectively, while ONT bulk had averages of ∼59.6% and ∼59.6% (Fig. 1I), likely owing to the usage of pychopper (see **Methods**). Single-cell long-read technologies had lower alignment rates overall, with ONT single-cell around ∼61.8% and ∼44.6%, and PB single-cell achieving ∼67.0% and ∼44.9%, each for genome and transcriptome alignment proportions, respectively.

In addition, for single-cell, we found that despite high raw read counts, PB and ONT only utilize 23.4% and 39.8% of raw reads respectively, once quantified with Oarfish (Fig. 1J). For ONT, the primary losses happen at the demultiplexing stage (38.9% of total reads discarded) and alignment stage (19.1% of total reads discarded), whereas PB loses most of its reads at the deduplication and alignment stages (53.9% and 16.0% of total reads discarded).

Overall, while all platforms and technologies identified the expected FMR1 transcriptional change, they also demonstrated variation in sequencing performance with regards to read quality, length, and efficiency.

### Quality control reveals platform- and technology-specific sequencing biases

Next, focused particularly on transcript detection and quantification differences between technologies and platforms.

Transcript detection, defined here as one count per million (CPM) or higher in all E3 replicates (long-read technologies) or one transcript per million (TPM) or higher in all E3 replicates (Illumina), when quantified against the GENCODE transcriptome (see **Methods**), differed greatly between platforms and technologies (Fig. 2A). Most notable were the long-read single-cell technologies, which detected the same ∼19,789 transcripts not detected in any bulk platform. While there were also transcripts that were detected only in the bulk technology, these were much rarer, pointing to potential quantification difficulties in the single-cell technology.

**Fig. 2:**
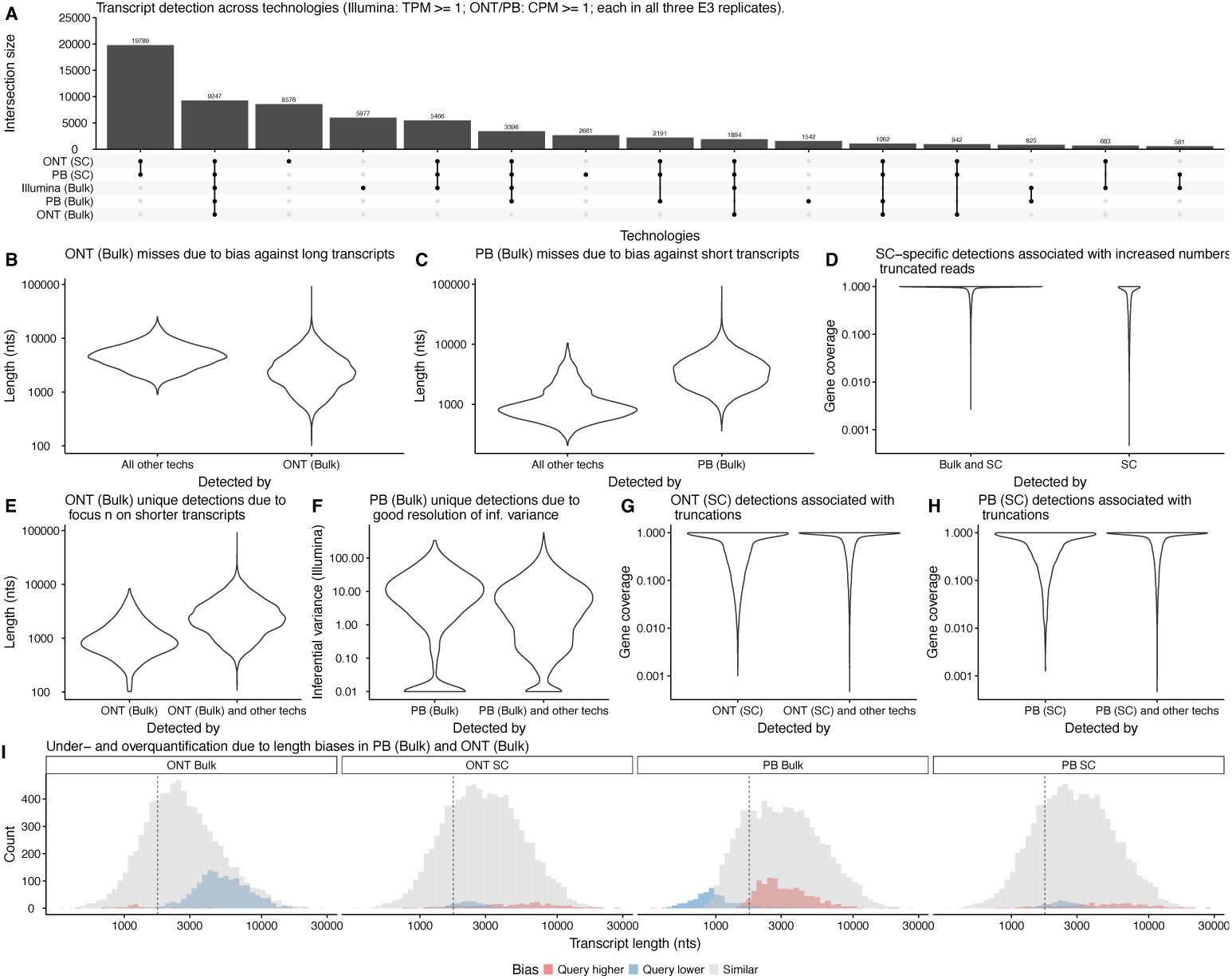
Biases of different platforms and technologies. **A.** Upset plot showing transcripts detected in all E3 samples across sequencing platforms and technologies. Single-cell platforms uniquely detect nearly 20,000 transcripts absent from any bulk dataset. **B.** Length distribution of transcripts detected in all technologies except ONT (Bulk) compared to transcripts detected in ONT (Bulk). ONT (Bulk) exhibits a clear bias against longer transcripts, yielding more shorter and fewer longer trancripts, compared to other technologies. **C.** Length distribution of transcripts detected in all technologies except PB (Bulk) compared to transcripts detected in PB (Bulk). PB (Bulk) exhibits a clear bias against shorter transcripts, yielding fewer shorter and more longer trancripts, compared to other technologies. **D.** Gene coverage (i.e., normalized exonic gene coverage proportion, per transcript, see **Methods**) of transcripts detected in at least one bulk technology and both single-cell long-read platforms compared to transcripts unique to single-cell long-read platforms. The single-cell long-read platforms exhibit a clear bias towards the detection of transcripts with incomplete gene coverage. **E.** Length distribution of transcripts detected uniquely in ONT (Bulk) compared to transcripts detected in ONT (Bulk) and at least one other technology or platform. ONT (Bulk) detects more transcripts that are very short compared to other technologies. **F.** Mean inferential relative variance of transcripts (see **Methods**) detected uniquely in PB (Bulk) compared to transcripts detected in PB (Bulk) and at least one other technology or platform. PB (Bulk) detects more transcripts that tend to have high inferential relative variance, pointing towards better resolution of transcripts belonging to complex genes. **G-H.** Gene coverage (i.e., normalized exonic gene coverage proportion, per transcript, see **Methods**) of transcripts detected uniquely in ONT (SC) (**G**) or PB (SC) (**H**), compared to detected in the corresponding long-read SC technology and at least one other technology. The single-cell long-read platforms again exhibit a clear bias towards the detection of transcripts with incomplete gene coverage, even compared to another single-cell long-read technology with the same library preparation (e.g., ONT single-cell compared to PB single-cell). **I.** Histogram of length of transcripts that are similar, or notably lower or higher in estimated abundance (see **Methods**) between a query technology and all other technologies, stratified by query technology, conditional on detection by all technologies. ONT (Bulk) and PB (Bulk) showed trends matching their detection properties, with ONT (Bulk) over-quantifying short transcripts and under-quantifying long transcripts, and PB (Bulk) showing the reverse behavior.

As such, we first investigated the biases of particular technologies that resulted in these technologies missing the detection of some transcripts. First, ONT bulk exhibited a clear bias against longer transcripts, starting around 5 kb (Fig. 2B). On the other hand, PB bulk showed the reverse bias, tending to miss transcripts that were shorter than 1.25 kb (Fig. 2C). Both of these are consistent with previous work (Wissel et al. 2026; You et al. 2025). Most of the other transcripts missed by one or both of the two bulk long-read technologies were specific to the single-cell long-read technologies. We found these transcripts to show a relatively large proportion of transcripts with low proportional gene coverages, suggesting that they may be caused by the lack of full-length transcripts in single-cell long-read technologies (Fig. 2D).

Next, we investigated why certain transcripts were only detected in some technologies or platforms. Transcripts that were uniquely detected in ONT bulk tended to be around 1 kb or shorter, matching its earlier bias against long transcripts (Fig. 2B, E). Transcripts that were uniquely detected with PB bulk demonstrated high inferential variance in Illumina bulk, suggesting that PB bulk may be especially good at resolving complex transcript structures (Fig. 2F). Lastly, transcripts that were uniquely detected by ONT single-cell or uniquely detected by PB single-cell exhibited a similar trend as transcripts common to both of the platforms. In particular, transcripts specific to single-cell lrRNA-seq technologies tended to have lower relative gene coverages, compared to transcripts common between single-cell lrRNA-seq and bulk technologies (Fig. 2H).

We then explored whether certain platforms would also under- or over-quantify certain groups of transcripts, in addition to the detection differences. Both ONT bulk and PB bulk showed quantification biases mirroring their detection biases, with ONT bulk over-quantifying (relative to all other technologies, see **Methods**) short transcripts and under-quantifying long transcripts (Fig. 2I). Similarly, PB bulk also mirrored its detection bias, under-quantifying short transcripts and over-quantifying long transcripts (Fig. 2I). ONT’s under-quantification of long transcripts and over-quantification of shorter transcripts likely reflects PCR-related inefficiencies, which we believe to be inherent to its chemistry. PB’s quantification bias is likely related to the number and stringency of size-selection steps typically performed during library preparation. Neither of the single-cell long-read technologies showed notable quantification biases (Fig. 2I).

While associated with gene coverage, we found that simple computational filters on the number of expressed transcripts within the gene (k) or the gene coverage of a detected transcript were unable to filter out transcripts solely detected in single-cell long-read quantifications while retaining other transcripts (Additional file 1: Supplementary Fig. 2A). Broadly speaking, the choice of quantification method did not affect any of the described biases notably (Additional file 1: Supplementary Fig. 2B-H). In addition, none of the technologies or platforms showed additional quantification biases based on gene coverage or inferential variance (Additional file 1: Supplementary Fig. 2I-J).

In summary, we caution researchers to carefully consider the choice of platform and technology depending on their research goals. We recommend that ONT bulk should be used if the interest is in shorter transcripts (less than 1.5 kb, or shorter) and PB bulk if most of the transcripts of interest are longer than 1.25 kb. In addition, we emphasize that single-cell long-read technologies using the ordinary 10X 3’ kit may not achieve a sufficient number of full-length reads for all transcript-level tasks, potentially leading to single-cell-specific transcript detections associated with truncations and other difficulties in transcript-level tasks such as DTE and DTU.

### Bulk quantification and spike-in analysis reveals the accuracy of Iscosceles, Miniquant, and Oarfish

We next explored quantitative lrRNA-seq performance, using both spike-in controls and GENCODE-mapped data, applying the most widely-used quantification tools: Bambu, Isoquant, Isosceles, Kallisto, Miniquant, and Oarfish (Chen et al. 2023; Prjibelski et al. 2023; Kabza et al. 2024; Loving et al. 2025; Zare Jousheghani et al. 2025; Li et al. 2025). To enable a balanced comparison across platforms, we downsampled the data to match the shallowest bulk sample, retaining 15M GENCODE reads, 75,000 SIRV reads, and 30,000 ERCC raw reads.

First, we benchmarked the memory usage and runtime performance of each method on the GENCODE data, using subsets of 1M, 2.5M, 5M, 10M, and 15M reads. For bulk quantification, Isoquant, Oarfish, and Kallisto demonstrated substantially lower memory usage compared to the other tools, with Kallisto showing near-constant memory consumption across all read depths. In contrast, runtime performance exhibited that Isoquant is comparatively slow, Oarfish is the fastest, followed by Miniquant, Kallisto, and Bambu (Fig. 3A–B).

**Fig. 3:**
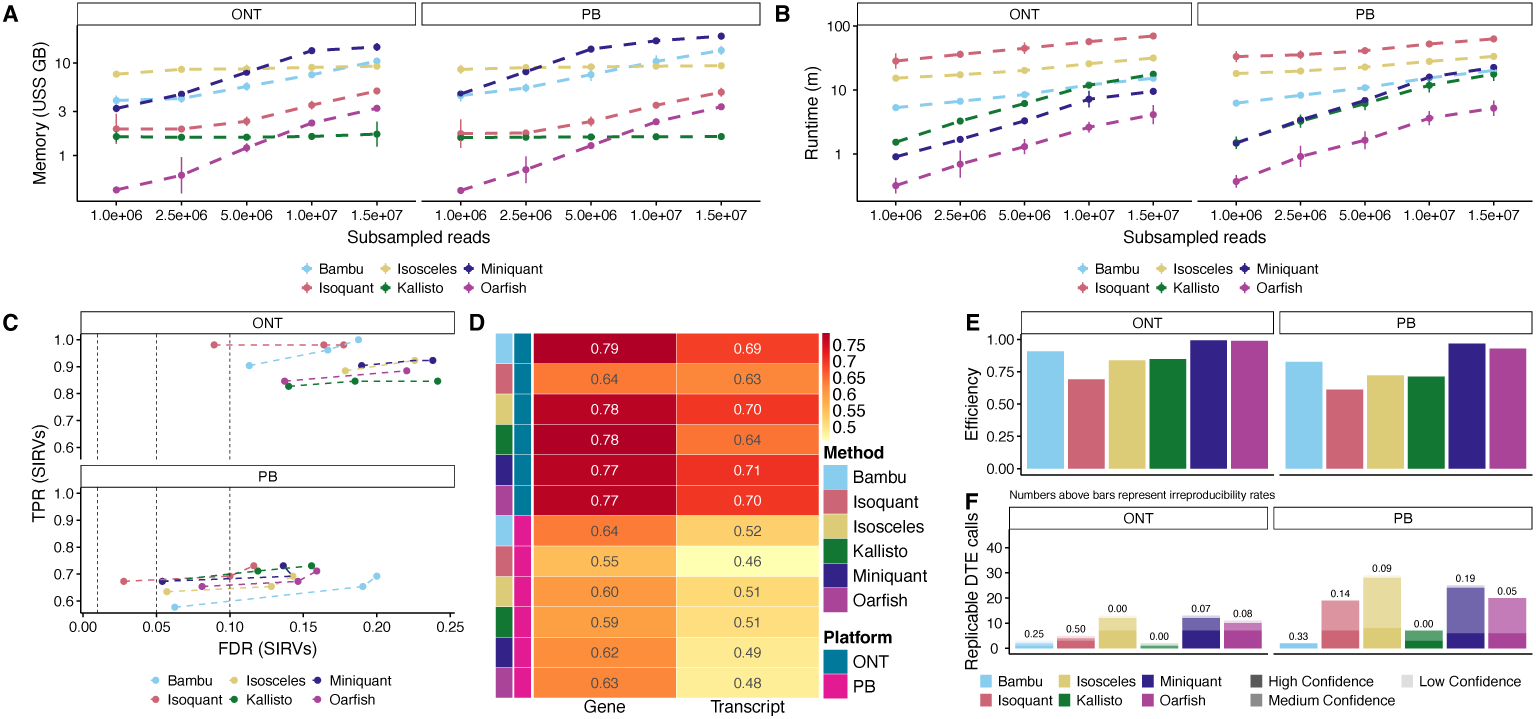
Bulk long-read quantification analysis. **A.** Memory usage of each quantification method on subsamples of 1M, 2.5M, 5M, 10M, and 15M reads. Error bars indicate variability across independent runs. **B.** Runtime of each quantification method on subsamples of 1M, 2.5M, 5M, 10M, and 15M reads. Error bars indicate variability across independent runs. **C.** DTE analysis on SIRVs, using the expected logFC as ground truth for validation. True positive rates (TPR) were computed based on agreement with expected logFC, while false discovery rates (FDR) were controlled in the analysis. For most methods and both platforms, TPR remains largely stable across modest variations in FDR. **D.** Spearman’s rank correlation between long-read sequencing platforms (log CPM, calculated using edgeR) and Illumina (logTPM, derived from Salmon) for both gene- and transcript-level GENCODE features, using features detected (1 or more CPM across all three E3 replicates for long-read technologies, and 1 or more TPM across all three E3 replicates for Illumina) in Illumina and at least one of the long-read platforms. Numbers in the table indicate the correlation coefficients. Across both platforms, Isosceles, Kallisto, and Oarfish exhibit the highest correlation to the filtered Illumina bulk quantifications. **E.** Quantification efficiency, defined as the ratio of subsampled reads to total counts, for each method and sequencing platform. Miniquant and Oarfish achieve the highest efficiency for both ONT and PB datasets. **F.** Replicable DTE calls for each quantification method across sequencing platforms, using FDR=0.01. Confidence is evaluated based on agreement across methods and platforms (see **Methods**). Isosceles, Miniquant, and Oarfish show the best balance between detection power and irreproducibility rate.

We next assessed quantification accuracy using the synthetic spike-in datasets. Specifically, we compared the observed versus expected concentrations of ERCC molecules, as well as the theoretical versus measured log fold change (logFC) for SIRVs. When stratifying transcripts by length (shorter or longer than 1.25kb), we again observed reduced performance for short transcripts in the PB data, whereas ONT achieved higher Spearman’s rank correlations with the true concentrations, not only for short transcripts but overall (Additional file 1: Supplementary Fig. 3A). When evaluating the accuracy of observed SIRV logFC stratified by transcript length, we again observed poor detection of transcripts shorter than 1.25kb in the PB data (Additional file 1: Supplementary Fig. 3B). When computing DTE results on the SIRVs, Isoquant achieved the highest true positive detection rate, while the other methods showed broadly comparable performance both in ONT and PB data (Fig. 3C).

After validating our spike-in data, we proceeded to benchmark the human GENCODE data. First, we computed the Spearman’s rank correlation between each long-read quantification method and Illumina data processed with Salmon using only E3 samples, and considering only genes or transcripts detected in Illumina and at least one quantification method of the long-read platforms in all E3 samples (one CPM or higher in all E3 samples for long-read technologies and one TPM or higher in all E3 samples for Illumina). In the human setting, expectation maximization (EM)-based methods, such as Isosceles, Kallisto, and Oarfish, achieved higher correlation with the Illumina data, and ONT, in general achieved higher correlations, likely due to PB strong length bias (Fig. 3D).

Finally, we evaluated the GENCODE data in terms of quantification efficiency and downstream tasks that rely on accurate read-to-transcript assignment (RTA), such as DTE and DTU. To assess quantification efficiency, we calculated the number of counts obtained per aligned read in each method. All methods, with the exception of Isoquant, showed comparable efficiency, with Oarfish and Miniquant performing best for both ONT and PB data (Fig. 3E).

For downstream analyses, no ground truth was available. We therefore constructed a reference set by leveraging concordance in both FDR and logFC across methods and datasets. Based on these two criteria, we stratified all transcript-level calls into three confidence tiers (see **Methods**).

For DTE, Miniquant and Isosceles identified the largest number of reproducible calls while maintaining an irreproducibility rate close to the nominal FDR of 0.01 for both PB and ONT (Fig. 3F, Additional file 1: Supplementary Fig. 3C). In contrast, Kallisto showed no irreproducible calls but identified fewer total calls across both PB and ONT datasets. Oarfish exhibited higher irreproducibility rates than Miniquant and Isosceles while calling slightly fewer reproducible calls in both PB and ONT, while Bambu called very few calls in general and exhibited high irreproducibility rates despite this (in ONT). Isoquant exhibited high irreproducibility rates and a low number of calls on ONT, but performed comparably to Oarfish on PB data.

For DTU, the comparison was more nuanced. Bambu, Oarfish, and Isoquant showed high irreproducibility rates in both ONT and PB data, although this also came with larger number of calls for Bambu and Oarfish but not Isoquant (Additional file 1: Supplementary Fig. 3D, E). Kallisto performed well on PB but had a high irreproducibility rate along with a very large number of calls on ONT data. Miniquant and Isosceles again performed well, not having any irreproducible calls in either ONT or PB data, although Miniquant had notably more calls than Isosceles. Overall, especially since there were few to no confident calls in all methods, we suggest viewing DTU results in conjuction with DTE results and preferring methods such as Isosceles and Miniquant, which perform more conservatively in terms of irreproducibility rates.

Overall, for the quantification of both ONT and PB bulk lrRNA-seq data, we recommend the usage of Isosceles, if sufficient computational resources are available, given its solid performance in spike-in data, close to calibrated irreproducibility rates, and among the highest numbers of reproducible calls. If computational resources are constrained, we recommend the usage of Miniquant or Oarfish, which produced overall very similar results, but resulted in either inflated irreproducibility rates (Oarfish) or a high number of low confidence reproducible calls (Miniquant) in DTU.

### Oarfish emerges as the leading method in single-cell quantification

To assess single-cell transcript-level quantification, we evaluated Bambu, Isosceles, Kallisto, and Oarfish, as these provided dedicated compatibility with lrRNA-seq single-cell data (Chen et al. 2023; Kabza et al. 2024; Loving et al. 2025; Zare Jousheghani et al. 2025). As with bulk, all samples were downsampled to match the shallowest sample (here, 60M reads; see **Methods**).

We computed the Spearman rank correlation between each long-read quantification method and Illumina bulk data, using only E3 samples and considering only genes or transcripts detected across all platforms in all E3 samples (one CPM or higher in all E3 samples for long-read technologies and one TPM or higher in all E3 samples for Illumina). To enable a direct quantitative comparison between the single-cell datasets and bulk Illumina sequencing, single-cell datasets were pseudobulked.

First, we evaluated the memory usage and runtime performance of single-cell lrRNA-seq quantification methods by subsampling datasets to 10M, 20M, 30M, and 60M reads. Kallisto and Oarfish consistently outperformed the other methods, showing nearly constant memory usage as the number of reads increased, with peak memory usage at 4.65GB and 3.17GB per sample, respectively. Notably, Oarfish also achieved the best runtime performance, maintaining an almost constant processing time across read depths, taking a maximum of 8 minutes to process one ONT sample and 3 minutes for one PB sample, whereas all other methods exhibited an approximately linear increase in runtime with the number of reads analyzed (Fig. 4A-B).

**Fig. 4:**
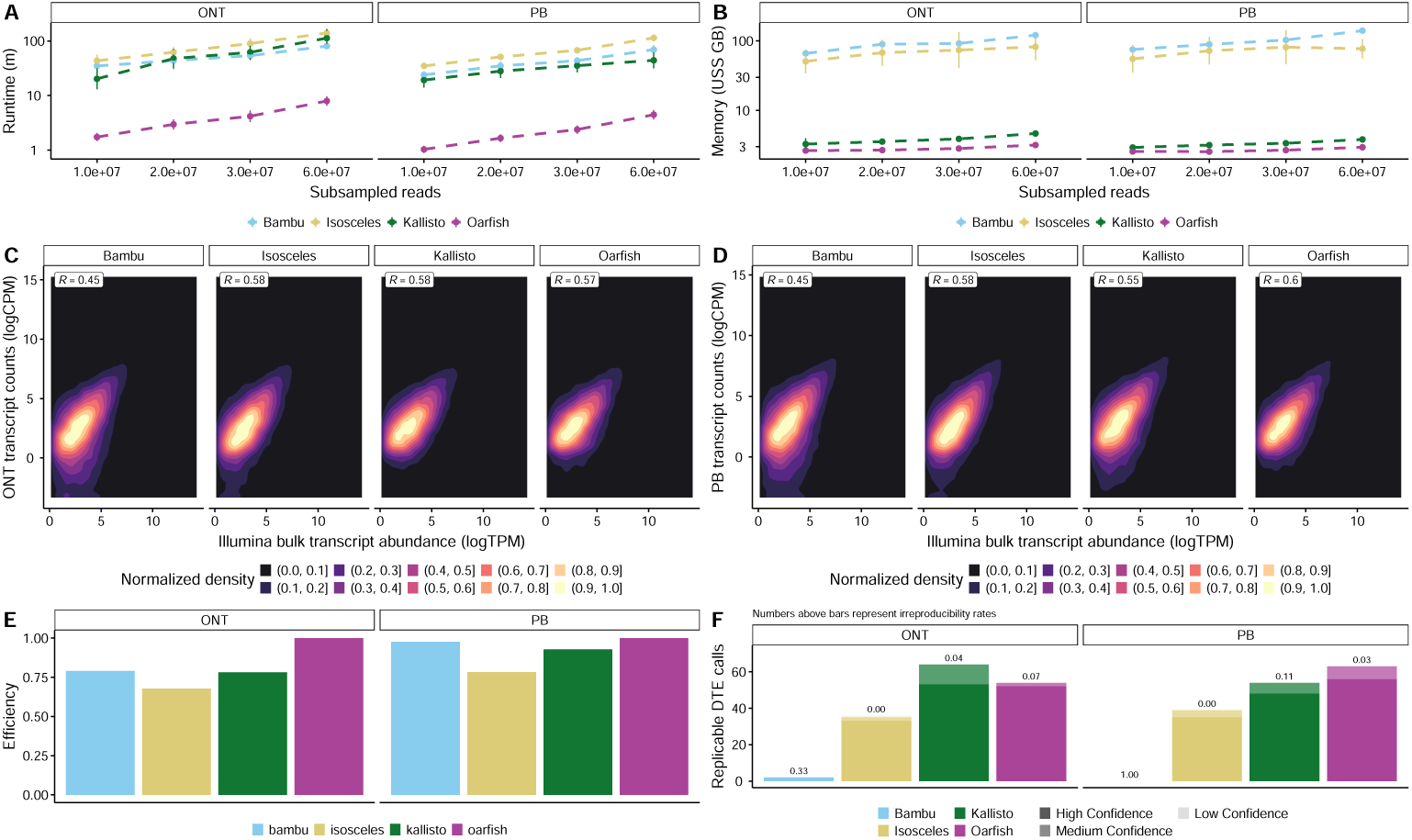
Single-cell datasets quantification analysis. **A.** Memory usage of each quantification method on subsamples of 10M, 20M, 30M, and 60M reads. Error bars indicate variability across independent runs. **B.** Runtime of each quantification method on subsamples of 10M, 20M, 30M, and 60M reads. Error bars indicate variability across independent runs. **C.** Spearman’s rank correlation between ONT pseudobulk (log CPM, calculated using edgeR) and Illumina bulk (log TPM, derived from Salmon) for transcript-level GENCODE features. Isosceles, Kallisto, and Oarfish perform comparably, while Bambu tends to underperform in terms of correlation to Illumina (Bulk). **D.** Spearman’s rank correlation between PB pseudobulk (log CPM, calculated using edgeR) and Illumina bulk (log TPM, derived from Salmon) for transcript-level GEN- CODE features. Isosceles, Kallisto and Oarfish perform comparably, while Bambu tends to underperform in terms of correlation to Illumina (Bulk). **E.** Quantification efficiency, defined as the ratio of subsampled mapped and deduplicated reads to total counts, for each method and sequencing platform. Oarfish achieves the highest efficiency for both ONT and PB datasets. **F.** Replicable DTE calls for each quantification method across sequencing platforms, using FDR=0.01. Confidence is evaluated based on agreement across methods and platforms (16). Oarfish shows the best balance between detection power and irreproducibility rate.

The single-cell transcript-level results highlighted once again the inherent difficulty of accurately quantifying human transcripts, with the maximum Spearman rank correlation of any single-cell lrRNA-seq quantification method of any technology only reaching 0.6 with the Illumina bulk data. Overall, for both ONT and PB single-cell, Isosceles, Kallisto, and Oarfish performed comparably when quantifying transcript expression, but Bambu underperformed compared to the other methods, reaching correlations around 0.45 for both technologies, compared to 0.55 to 0.60 for other methods (Fig. 4C-D).

Lastly, we evaluated the quantification efficiency of different methods, along with their performance in downstream tasks such as DTE, and DTU. Quantification efficiency was assessed by calculating the number of counts obtained per aligned and de-duplicated read for each method (see **Methods**). All methods showed comparable efficiency, with Oarfish performing best for both ONT and PB data (Fig. 4E). For downstream analyses, we applied the same strategy used in Fig. 3 to establish the ground truth for the GENCODE data, albeit with slightly more restrictive criteria (see **Methods**). Here, Oarfish single-cell quantification mode outperformed the other methods in terms of power, while showing a similar irreproducibility rate, although the irreproducibility rate for single-cell DTE was generally heavily inflated for most methods (Fig. 4F, Additional file 1: Supplementary Fig. 4A).

Overall, we thus recommend the usage of Oarfish for quantifying both ONT and PB single-cell lrRNA-seq data, given its excellent computational performance (which is a greater bottleneck for single-cell), and a high number of reproducible DTE calls at moderate irreproducibility rates. If sufficient computational resources are available, we also recommend the usage of Isosceles, especially if conservative calls are desired, given that the method achieved moderate to high reproducible calls while maintaining a near-calibrated irreproducibility rate for both DTE and DTU.

### PB single-cell data requires approximately three to four times as high depth as bulk data

When designing lrRNA-seq experiments utilizing both bulk and single-cell formats, it is useful to establish read depths that lead to equivalent downstream results between technologies or platforms. For this, we ran DTE, establishing reproducible and irreproducible calls as previously (see **Methods**; briefly, reproducible calls needed to be shared with at least one other quantification method in a different platform or technology; irreproducible calls were calls on transcripts that were detected in other platforms or technologies that disagreed notably in magnitude or directionality) and calculated how many reads corresponded to the same number of calls in another platform or technology.

When considering DTE, ONT bulk called fewer replicable DTEs than other bulk technologies, which led to an almost 1:1 ratio when considering depth equivalence between ONT bulk and single-cell, with 10M ONT bulk reads corresponding to approximately 11.4M ONT single-cell reads and 15M ONT bulk reads corresponding to roughly 17.1M ONT single-cell reads (Fig. 5A, top left). PB’s depth equivalence was more nuanced, with 5M PB bulk reads corresponding to 13.3M PB single-cell reads, but 10M bulk reads to 35.3M single-cell reads and 15M to 40.6M, leading to an average ratio of 3.12 for the two larger depth comparisons (Fig. 5A, top right).

**Fig. 5:**
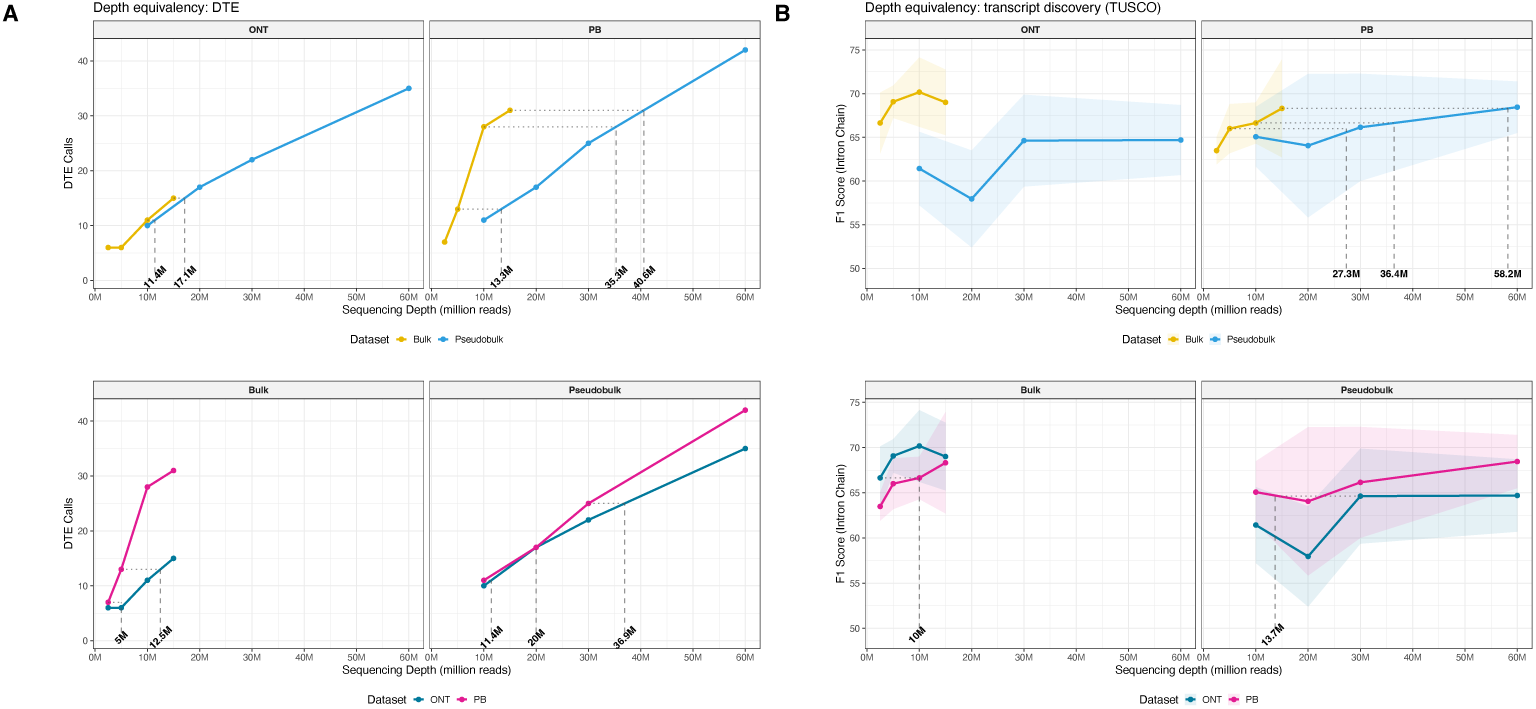
Downstream tasks for cross-platform evaluation. **A.** Numbers of DTE detected at varying subsampled depths using Isosceles counts (bulk) and Oarfish counts (single-cell). Bold numbers highlight empirical equivalences between datasets or technologies (e.g., approximately 31.8M PB single-cell reads are required to recover the same number of DTEs detected with 10M PB bulk reads). Top: Bulk vs pseudobulk. Bottom: PB vs ONT. **B.** F1 score of rediscovering the universal TUSCO gene set at different downsampling depths in multiple platforms and technologies. Bold numbers highlight empirical equivalences between datasets or technologies (e.g., approximately 36.4M PB single-cell reads are required to recover the same number of transcripts discovered with 10M PB bulk reads). Top: Bulk vs pseudobulk. Bottom: PB vs ONT.

Next, we compared different technologies, for the same platform. Here, ONT bulk once again underperformed due to its fewer replicable DTE calls, with 2.5M PB bulk reads corresponding to 5M ONT bulk reads, and 5M PB corresponding to 12.5M ONT reads, yielding an average ratio of 2.25 (Fig. 5A, bottom left). We note that we do not necessarily believe this number to be representative, as it is heavily driven by ONT and PB bulk’s preference towards specific transcript lengths (Fig. 2B, I). Between single-cell technologies, PB had a slight edge in terms of DTE calls, with 10M PB reads corresponding to 11.4M ONT reads, and 20M, 30M corresponding to 20M and 36.9M, respectively, giving an average ratio of 1.12, indicating approximately equal performance (Fig. 5A, bottom right).

Next, we investigated transcript discovery, focusing on rediscovering the TUSCO set as a pseudo-ground truth (Liu et al. 2025), and the “rediscovery” of specific GENCODE transcripts (see **Methods**). We used Bambu for transcript discovery in all considered platforms and technologies, with a consistent NDR (see **Methods**). For TUSCO, with the exception of the platform comparison for PB, we found consistent differences between both platforms and technologies, which made the establishment of depth equivalences impossible (Fig. 5B). Overall, ONT bulk consistently outperformed ONT pseudobulk (Fig. 5B, top left). PB data was largely congruent with DTE results, with bulk data consistently requiring notably fewer reads, with 5M bulk reads corresponding to 27.3M single-cell reads, 10M to 36.4M, and 15M to 58.2M, yielding an average ratio of 4.33 (Fig. 5B, top right). ONT bulk outperformed PB bulk (Fig. 5, bottom left), and PB single-cell outperformed ONT single-cell (Fig. 5, bottom right). Lastly, the GENCODE re-discovery did not allow for the establishment of any depth equivalencies, with both bulk technologies far outperforming the corresponding single-cell platform (Additional file 1: Supplementary Fig. 5A, left). Between bulk technologies, PB outperformed ONT, and the two single-cell technologies performed broadly similar (Additional file 1: Supplementary Fig. 5A, right). Interestingly, the bulk technologies showed decreasing performance at higher depth, despite a constant NDR given to Bambu (see **Methods**), which may nevertheless be related to filtering. Overall, given the lack of overlap in performance for transcript discovery and the large differences in ONT bulk quantifications to other platforms and technologies, depth equivalency between ONT bulk and single-cell technologies remains unclear. Similarly, we do not think depth equivalencies can be robustly established between ONT bulk and PB bulk, given the impact of the length biases between the two methods (Fig. 2B, E, I). For the comparison between PB bulk and PB single-cell, however, the ratio was, on average, 3.12 for DTE and 4.33 for transcript discovery. Thus, taken together, PB single-cell sequencing requires approximately three- to four-fold greater depth than bulk to achieve comparable performance in both transcript discovery and differential transcript expression, which seems largely attributable to its lower proportion of exonic reads and PCR duplication yielding a greater drop from raw reads to counts (Fig. 1I-J). The comparison between the two single-cell technologies also yielded usable results, indicating that the two technologies are largely comparable, with PB needing slightly fewer reads to achieve a similar number of DTE calls, and it being largely unclear for discovery (given PB’s slight outperformance for TUSCO and ONT’s for GENCODE).

Overall, we thus recommend budgeting for higher targeted read depth in the following ways. If comparable results to PB bulk data should be achieved, single-cell PB should be sequenced at approximately 3-4 times the targeted depth. If comparable results to PB single-cell data should be achieved, single-cell ONT should be sequenced at approximately the same depth.

### Case study: Matching long-read RNA-seq experimental design to the corresponding research question is crucial

Our study argues for the importance of considering sequencing technology and platform, quantification method, and read depth when designing lrRNA-seq experiments. In particular, making suboptimal choices in one or more of these makes specific research questions notably more difficult.

To illustrate this, we highlight an example demonstrating that even genes of low to moderate complexity can, at times, be poorly quantified using single-cell longread data which especially affects differential splicing analyses. As an exemplary case, we examined the *WASF3* gene, which displayed markedly different count-proportion profiles between bulk and single-cell samples across both long-read platforms. We compared the read-level evidence in both bulk and single-cell data for ONT and PB, and observed discrepancies in the transcript proportions despite strong read support and a limited transcript repertoire (Fig. 6A-C). These observations are potentially attributable to a combination of reverse transcription truncation and internal priming, which can be especially problematic for single-cell long-read libraries, due to the lower proportion (or even lack of) full-length reads.

**Fig. 6:**
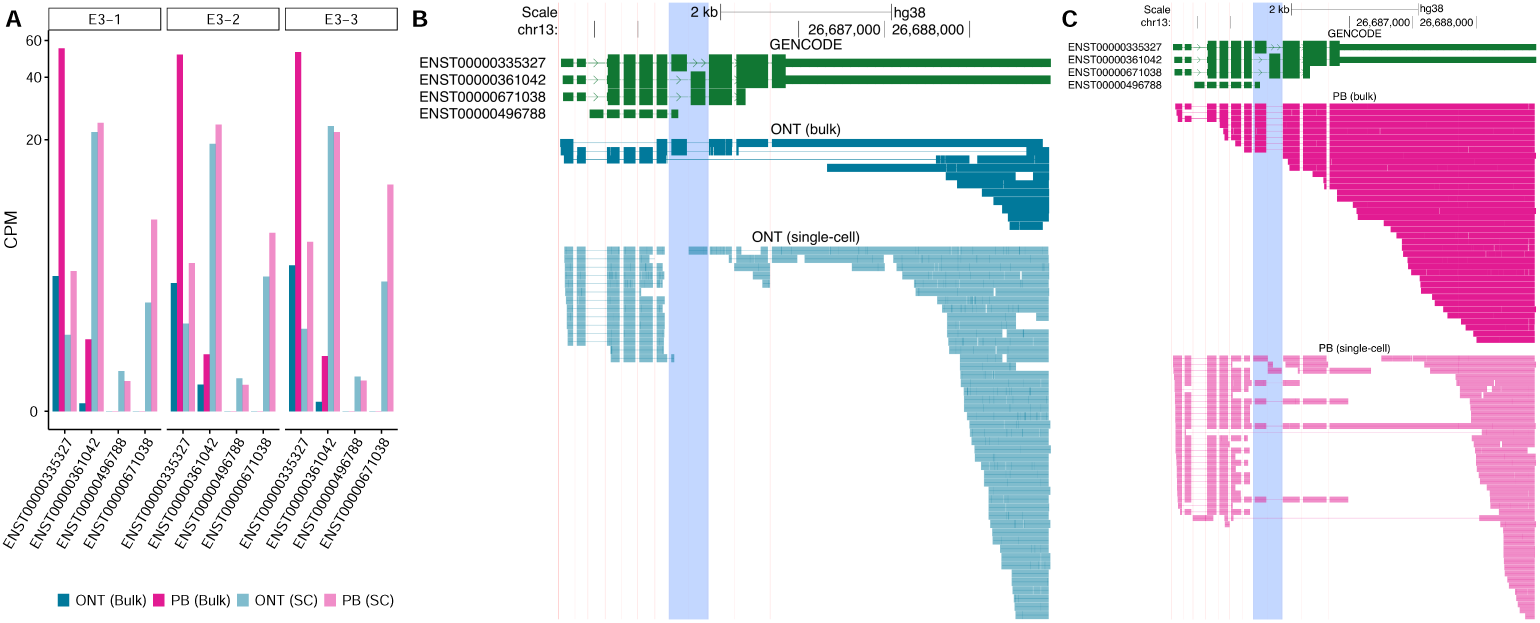
Case study illustrates challenges in read-to-transcript assignment within single-cell long-read technologies. **A.** CPM values for the gene *WASF3* across long-read platforms and technologies for all three E3 replicates. In pseudobulk datasets, counts are dispersed across multiple *WASF3* transcripts, whereas in bulk datasets expression is dominated by a single transcript, ENST00000335327. **B.** Genome browser track of expressed GENCODE V45 transcripts for the *WASF3* gene, along with genome alignments of the ONT bulk and ONT single-cell E3-1 samples. **C.** Genome browser track of expressed GENCODE V45 transcripts for the *WASF3* gene, along with genome alignments of the PB bulk and PB single-cell E3-1 samples.

Overall, our case study shows one example of the importance of aligning (in this case) the choice of platform and sequencing technology with the research goal in question.

## Discussion

Our evaluation of lrRNA-seq across platforms and technologies was conducted in a human neuronal transcriptome, a system, to the best of our knowledge, not previously used in cross-platform long-read studies, highlights the necessary considerations and trade-offs among sequencing depth, platform selection, technology format, and quantification algorithm choice.

In our study, we have evaluated lrRNA-seq platforms across both bulk and single-cell formats and quantification methods with the aim of highlighting key performance metrics related to transcript-level analyses. We observed distinct technical biases for different sequencing platforms and technologies. PB bulk struggled with the detection and quantification of transcripts shorter than 1.25 kb, likely due to its size selection. Conversely, ONT bulk data captured shorter transcripts effectively but had issues with transcripts longer than 5 kb, in both detection and quantification. We observed large differences in detected transcripts in single-cell data that did not appear in matched bulk datasets, for both PB and ONT single-cell datasets. Our read-level analysis of genes, such as WASF3, suggests that this may be largely technical rather than biological. Single-cell protocols, such as 10X, can promote internal priming and reverse-transcription truncation, generating fragments that quantification algorithms struggle to assign accurately to a single transcript, similar to srRNA-seq. The WASF3 gene in particular demonstrated substantial read truncation within the fifth exon in single-cell data. This pattern is also consistent with a known difference in reverse transcription efficiency between bulk and single-cell protocols. In droplet-based single-cell workflows, reverse transcription occurs in a sub-optimal biochemical environment, including the presence of cellular lysate, lipids, proteins, and partitioning reagents, which can impair enzyme performance and increase premature termination. By contrast, bulk cDNA synthesis typically utilizes pristine extracted RNA and optimized reaction conditions. As a result, single-cell libraries frequently yield shorter, more truncated cDNAs, which can be misinterpreted by quantification algorithms as distinct transcript isoforms, inflating transcript counts and distorting proportion estimates, which makes DTE and DTU analyses more challenging.

Some computational methods for transcript quantification of lrRNA-seq performed notably better than others in our benchmarks. While simpler tools like Isoquant performed well on synthetic spike-ins (SIRVs and ERCCs), they struggled to resolve the complex realities of the human transcriptome. Specifically, Oarfish and Isosceles consistently outperformed others in accuracy and downstream power, while maintaining decent computational efficiency. Overall, we recommend Oarfish for single-cell applications and Isosceles for bulk analyses, for both ONT and PB.

Finally, our downsampling analyses underscore that single-cell datasets commonly require depths on the order of three to four times as large as comparable bulk lrRNA-seq datasets. It is likely that similar depth equivalency recommendations are advisable between ONT bulk and single-cell, but we were not able to directly establish them in our work.

Of course, our study has several limitations. Most notably, lrRNA-seq methods and technologies are constantly advancing. As such, our work represents only a snapshot in time and should not be expected to necessarily generalize to future technologies, library preparation methods or datasets. In addition, our analyses of single-cell lrRNA-seq methods and technologies were focused exclusively on libraries prepared using the 10X 3’ kit, and performance may not be representative of other single-cell lrRNA-seq platforms. Furthermore, since our model does not provide a robust ground-truth for endogenous transcripts (that is, aside from the SIRV spike-ins), we rely on cross-platform and/or cross-method comparisons to calculate reproducible and irreproducible calls for downstream tasks such as DTE and DTU, which is less robust than gold standard ground truths. Also, the SIRV spike-ins we utilize are known to not be perfectly representative of human or other mammal transcriptomes in terms of length, splicing patterns, and other aspects, which may make our spike-in analyses less valuable.

Despite these limitations, we believe our work conveys valuable points about the current landscape of technologies and methods for bulk and single-cell lrRNA-seq. In addition to this, it may also serve as a useful framework to enable future bench-marking work. Furthermore, in any case, our dataset should serve as a valuable comparison platform for researchers interested either in FMR1 biology or lrRNA-seq benchmarking.

Overall, there are several aspects of our work that align with the efforts of previous contributions. In particular, Scoones et al. (2025) previously highlighted that 10X Chromium-based single-cell lrRNA-seq approaches frequently resulted in very short, potentially truncated transcripts, which is mirrored in both our case study and the large number of potentially spurious transcript detections by both single-cell ONT and PB. In addition, the authors highlighted the high drop from raw to quantifiable reads, which we also showcase in our quality control. Several of the biases found in our study were also mirrored by the results of You et al. (2025). In particular, the bias against long transcripts in ONT bulk and the bias of PB bulk against short transcripts were both also found by You et al. (2025), the latter in addition also by Wissel et al. (2026). Despite this, there are several aspects of our work that, to the best of our knowledge, were not previously reported or considered. First, we highlight that ONT bulk detects several shorter transcripts that tend to be missed by other technologies, while PB bulk seems to detect more transcripts that exhibit high inferential variance in srRNA-seq. Furthermore, to the best of our knowledge, our work is the first to evaluate different single-cell lrRNA-seq quantification methods. Similarly, while the dropoff in single-cell lrRNA-seq read counts relative to raw counts was previously shown by Scoones et al. (2025), we are, to the best of our knowledge, the first to construct raw read-depth equivalencies between bulk and single-cell, supporting more informed decisions about experimental design. Lastly, we believe that our single-cell lrRNA-seq dataset provides a unique combination current library preparation chemistries, multiple conditions with replication, and deep sequencing depth (approximately 100 M) to enable differential expression and other downstream tasks.

In summary, while lrRNA-seq provides an unprecedented view of the transcriptome, neither the technologies, platforms, nor quantification methods for it are free of biases. Thus, we have provided a comprehensive set of recommendations regarding sequencing platform, depth, technology format, and quantification methods, which we hope will guide researchers aiming to resolve complex transcript architectures using lrRNA-seq.

Although this study utilized a neuronal model of Fragile X Syndrome, the design principles and platform-specific constraints identified here are likely applicable to transcriptomic studies of other complex tissues.

## Authors’ contributions

G.B. and D.W. contributed equally to this work.

G.B. and D.W. performed all computational analyses, generated the figures, conducted the statistical analyses, and conceived the study with input from S.H., W.S., M.D.R., and R.S. W.S., M.D.R., and R.S. supervised the work, and M.D.R. and R.S. acquired funding and resources for the study. A.P. contributed to computational analyses of the single-cell lrRNA-seq data. T.C. designed and supervised the neuronal differentiation strategy. K.R., S.P., J.R.C., K.S., and Z.M.N. performed the CRISPR-mediated genome editing, conducted neuronal differentiation experiments, carried out the 10x Genomics library preparation for single-cell RNA sequencing, and performed the Illumina single-cell RNA-seq experiments. A.B. and W.S. performed library preparation and sequencing for ONT samples. C.N., and Z.M. provided sequencing platform expertise and resources. S.J.C. provided biological resources and supervision. S.H. contributed formal analysis support. The manuscript was written by G.B. and D.W., and was supervised and edited by M.D.R., W.S., and R.S. The manuscript was reviewed and approved by all authors.

## Methods

### Generation and characterization of FXS-patient-derived iPSC lines

The FXS patient-derived iPSC line WC005i-FX11-7 was obtained from WARF/Wi-Cell. This line carries a full mutation (*>*435 CGG repeats) in the 5’ untranslated region (UTR) of the *FMR1* gene, which results in its transcriptional silencing (Doers et al. 2014). To facilitate the detection of endogenous *FMR1* reactivation, the iPCSs were CRISPR-modified to introduce a reporter cassette (P2A-GFP-T2A-BlastR) into the 3’-end endogenous *FMR1* locus, in-frame with the coding sequence. Upon clonal selection of the FXS reporter line, clone E3 (hereafter FXS E3), an isogenic corrected (rescue) line was generated. The CGG repeat expansion was excised using CRISPR-dependent deletion and sgRNA pair from Xie et al. (2016), spanning the CGG repeats. Reactivation of the FMR1 locus was verified via fluorescent imaging (GFP), expression profiling, and WES assay using anti-FMRP antibodies (Cell Signaling, cat. No. 4317).

### FXS NGN2-iNeurons Differentiation

Glutamatergic NGN2-induced neurons (NGN2-iNeurons) were generated from Fragile X Syndrome (FXS) induced pluripotent stem cells (iPSCs) over a 21-day period using a doxycycline-inducible NGN2 cassette activation adapted from Peitz et al. (2020). Briefly, following the induction and Day 2 passaging onto polyornithine/laminin-coated plates, cells were cultured in DAPT-supplemented neuronal medium without AraC. Neuron maintenance from Day 7 involved thrice-weekly half-media changes containing BDNF and laminin to sustain final concentrations of 1:1000 and 1:500, respectively. At Day 21, cells were collected for downstream single-cell and bulk (SR/LR) transcriptomic profiling.

### RNA extraction and spike-in injection

Total RNA was extracted from NGN2-iNeurons using TriZol. Sample QC on total RNA was performed using Qubit RNA High Sensitivity (Thermo Fisher Scientific, Q32852) to measure concentration and RNA ScreenTape Analysis (Agilent, 5067-5576) to assess RNA integrity. All RNA samples exhibited RIN scores 9.4. ERCC RNA Spike-In Mix (Thermo Fisher Scientific, 4456740) was added to each sample to comprise an estimated 0.5% of the mRNA. Additionally, SIRV Spike-In controls (Lexogen, 025.03), (either set E0 or E2) were added to the samples to comprise 0.5% of the mRNA.

### 10X Genomics single-cell cDNA and library generation

Single-cell RNA-seq was performed using the 10X Genomics Chromium instrument with the NextGEM Single-cell 3’ Kit v3.1. Viability of iNeurons was assessed prior to injection and exceeded 95% in all replicates. Approximately 6,000 cells were targeted for each injection/replicate. After the Post-GEM-RT Cleanup & cDNA Amplification (Step 2), cDNA was stored for downstream short- and long-read library preparation and sequencing.

### Library preparation and sequencing

For the Illumina bulk dataset, 250ng of total RNA from each replicate was used as input for TruSeq Stranded mRNA library preparation kit (Illumina, 20020595). Samples were sequenced on an Illumina NovaSeq X Plus using PE 150bp reads and targeting approximately 50M reads per sample. For the PB bulk dataset, 300 ng of total RNA with a RIN greater than 7 was used as input. Libraries were prepared using the Iso-Seq Express 2.0 Kit (PB, 103-071-500) together with the Kinnex PCR 8-fold Kit (PB, 103-071-600) following the manufacturer’s instructions. cDNA amplification was performed for ten cycles, and the resulting libraries were purified using SMRTbell Cleanup Beads (PB, 102-158-300). Library concentration and fragment size were confirmed with the Agilent Tapestation 4200 and Qubit Flex Fluorometer prior to sequencing. Sequencing was performed on the PB Revio platform using the Revio Polymerase Kit (102-817-600) with diffusion loading and a 24-hour movie time per SMRT Cell. For the bulk ONT dataset, 500ng of total RNA (with spike-ins) was prepared using the cDNA-PCR Sequencing V14 kit (Oxford Nanopore Technologies, SQK-PCS114.24). 14 cycles of PCR amplification were performed. Libraries were sequenced using PromethION flow cells (FLO-PRO114M) on a PromethION 24 instrument. 3 samples were multiplexed per flowcell using the dorado dnȧr10.4.1̇e8.2̇400bpṡhac@v4.3.0 basecaller. For the Illumina single-cell dataset, library construction was performed using fragmentation, end repair, A-tailing, and adaptor ligation. Final libraries were generated via PCR amplification using the Dual Index Kit TT Set A to attach unique i5 and i7 indices. Library size distribution and concentration were assessed using an Agilent TapeStation D1000 tape and Qubit fluorometer, respectively, prior to sequencing on an Illumina NovaSeq 6000 Patterned. For the PB single-cell dataset, amplified cDNA generated after Step 2 (Post-GEM- RT Cleanup and cDNA Amplification) from the 10x Genomics Chromium Next GEM Single-Cell 3’ Kit v3.1 workflow was used as input for PB library construction. Two sequencing rounds were performed using distinct library preparation chemistries to evaluate consistency and performance across versions of the PB single-cell RNA workflow.

In the first round, 75 ng of amplified cDNA was processed using the MAS-Seq for 10x 3’ Concatenation Kit (PB, 102-407-900). This protocol enables the concatenation of multiple short 3’ fragments into long molecules while preserving 10x cell barcodes and unique molecular identifiers (UMIs). SMRTbell libraries were cleaned with SMRTbell Cleanup Beads (PB, 102-158-300) and amplified for nine cycles during cDNA synthesis. Sequencing was performed on the PB Revio instrument using the Revio Polymerase Kit (102-817-600) with diffusion loading and a 24-hour movie time per SMRT Cell. Each run generated high-fidelity (HiFi) reads. Library quality and concentration were confirmed using the Agilent Tapestation 4200 and Qubit Flex Fluorometer prior to sequencing. In the second round, 75 ng of amplified single-cell cDNA was prepared using the Kinnex Single-Cell RNA Kit (PB, 103-072-200) following the manufacturer’s Iso-Seq workflow for single-cell 3’ libraries. The protocol included nine amplification cycles and cleanup with SMRTbell Cleanup Beads (PB, 102-158-300). Sequencing was again performed on the PB Revio using the Revio Polymerase Kit (102-817-600), diffusion loading, and 24-hour movies per SMRT Cell.

For the ONT single-cell dataset, 10ng of cDNA amplicons prepared using 10X Genomics (NextGEM Single Cell 3’ Kit v3.1) were processed according to the single-cell transcriptomics protocol (SSṪv9198̇v114̇revĖ06Dec2023) from Oxford Nanopore Technologies. cDNA was amplified using the standard primers. Pull-down of biotin-tagged cDNA was performed with M280 magnetic streptavidin coated beads (Invitrogen, 11205D) prior to a second round of PCR using PRM primers (Oxford Nanopore Technologies, EXP-PCA001). Sample QC was performed using both Qubit dsDNA High-sensitivity (Thermo Fisher Scientific, Q32854) and D5000 ScreenTape Assay (Agilent, 5067-5588). Finally, each sample of amplified cDNA was prepared using the Ligation Sequencing Kit V14 (SQK-LSK114) and prepared for loading on a single PromethION flow cell (FLO-PRO114M) prior to sequencing with the PromethION 24 instrument, using the dorado dnȧr10.4.1̇e8.2̇400bpṡhac@v4.3.0 basecaller.

### References

For human data, we used GRCh38 as the reference genome (GCȦ000001405.15̇GRCh38̇nȯalṫanalysiṡset.fna.gz) and the primary assembly of GENCODE V45 as the reference transcriptome. For the SIRV and ERCC spike-ins we used the reference genome and transcriptome provided by Lexogen for SIRV set four, which contains all SIRVs along with ERCCs.

### Bulk RNA-seq preprocessing

Illumina samples were preprocessed using triṁgalore 0.6.10 to trim for adapters and quality. Samples were then aligned to a joint genome containing both the human and spike-in genomes using STAR 2.7.11b, using default parameters and outputting both genome and transcriptome alignments (–quantMode TranscriptomeSAM). Aligned reads were then divided into GENCODE, SIRV, and ERCC reads. PB samples were preprocessed by running ccs 8.0.0, pbtrim 1.0.0, jasmine 2.2.0, and lima 2.9.0. Following this, we ran skera-split 1.2.0 for S-read generation, lima 2.13.0 for primer removal, and finally isoseq refine 4.3.0 for FLNC read generation. Samples were then aligned to a joint genome containing both the human and spike-in genomes using minimap2 2.28-0, using standard parameters for spliced alignment with accurate PB long reads (-ax splice:hq -uf) (Li 2018). Aligned reads were then divided into GENCODE, SIRV, and ERCC reads. Additionally, SIRV and ERCC reads were realigned to the genome using minimap2 with parameters optimized for spike-ins (all parameters as before, except adding –splice-flank=no). Each category of reads was also aligned to the transcriptome using minimap2 2.28-0 with optimal parameters for PB HiFi reads (-ax map-hifi). ONT samples were preprocessed using pychopper 2.7.10 for kit V14 (-k PCB114) to remove primers and filter for full-length reads. Samples were then aligned to a joint genome containing both the human and spike-in genomes using minimap2 2.28-0, using the standard parameters for spliced alignment with long reads (-ax splice -uf). Aligned reads were then divided into GENCODE, SIRV and ERCC reads. Additionally, SIRV and ERCC reads were realigned to the genome using minimap2 with parameters optimized for spike-ins (all parameters as before, except adding –splice-flank=no). Each category of reads was also aligned to the transcriptome using minimap2 2.28-0 with optimal parameters for ONT reads (-ax map-ont).

### Single-cell RNA-seq preprocessing

Illumina samples coming from different lanes were concatenated together. PB samples were preprocessed by running ccs 8.0.0, pbtrim 1.0.0, jasmine 2.2.0, and lima 2.9.0. Following this, we ran skera-split 1.2.0 for S-read generation, lima 2.13.0 for primer removal, isoseq tag for barcodes and UMIs detection, isoseq refine for FLNC read generation, isoseq correct to correct barcodes, and finally isoseq groupdedup for PCR deduplication. All the isoseq commands were run using version 4.3.0. Reads were aligned using minimap2 2.28-0 to both the genome and the transcriptome. Genome alignments were performed with the standard parameters for spliced long-read alignment (-ax splice -uf), while transcriptome alignments used the recommended settings for PB HiFi reads (-ax map-hifi). ONT samples were preprocessed by running blaze 2.5.1 to demultiplex barcodes and UMIs. Following this step, samples were aligned to the transcriptome using minimap2 2.28-0, using standard parameters for ONT reads (ax map-ont), and deduplication was accomplished by filtering primary alignments and running umi̇tools dedup 1.1.6. Before proceeding, secondary alignments of the deduplicated reads were restored. Reads were also aligned to the genome using minimap2 2.28-0 with standard parameters for spliced alignment with long reads (-ax splice -uf).

### Read statistics and FMR1 rescue validation

High-level QC was performed across all datasets, independent of quantification methods. We first assessed the total number of reads, raw and aligning to the human, ERCC, or SIRV genomes, respectively. Then, to validate the successful rescue of FMR1 expression, we performed DGE on each platform and technology, only considering genes with at least one CPM (long-read technologies) or TPM (Illumina) in at least three samples and tested for DGE using edgeR’s glmQLFit 4.6.3. We computed MDS dimensionality reductions to assess sample clusterings. MDS was computed using edgeR 4.6.3 with default parameters on transcript expression values. Lastly, we evaluated replicability on transcript expression values, considering only transcripts that had at least one CPM in at least three samples (long-read technologies) or at least one TPM in at least three samples (Illumina).

### Quality control and quantification biases

We evaluated the raw read length, base qualities, empirical error rates, and the number of covered junctions, each on 1M randomly sampled reads aligning to the human genome for all considered platforms. In addition, the 3’ bias of all platform-technology pairs was evaluated on 2,500 randomly sampled GENCODE transcripts, within four transcript size bins, less than one kb, between one and two kbs, between two and three kbs, and between three and six kbs. The empirical SNV rate was calculated by parsing the BAM MD tag, and the indel rate was calculated by parsing the CIGAR string using pysam 0.23.0. Covered junctions were calculated as the number of introns of length 20 or longer in the cigar string of the genome alignments of each platform-technology pair using pysam 0.23.0. 3’ bias was computed by RSeQC 5.0.4.

For quantification biases and general detection, we considered detection as all E3 samples having one CPM (long-read technologies) or TPM (Illumina) or higher for a particular transcript. Numbers at each step of the single-cell processing pipelines for ONT and PB were calculated manually using samtools 1.22.1 and cached for plotting. Inferential relative variance was computed using computeInfRV using fishpond 2.14.0.

### Bulk quantification and spike-in analysis

Within-platform comparisons were performed using downsampled read sets, ensuring that each quantification method for a given platform received an identical subset of reads for fair evaluation. Read depths were standardized to 15M reads for GENCODE data, 75 thousand reads for SIRVs, and 30 thousand reads for ERCCs. These values were determined based on the shallowest sample available across the three platforms. Quantification for ONT and PB was performed using Isoquant, Bambu, Oarfish, Kallisto, Isosceles and Miniquant. Isoquant 3.6.3 was run on genome alignments of both platforms with –polyȧrequirement never and adjusting –datȧtype per platform. Bambu 3.4.0 was run on genome alignments of both platforms with default parameters. Oarfish 0.6.3 was run on transcriptome alignments of both platforms with model coverage applied (–model-coverage), no filters and a bin width of 100. Kallisto 0.51.1 was run on raw FASTQ reads of both platforms by adjusting the parameters for each platform and otherwise default parameters. Isosceles 0.2.1 was run on genome alignments of both platforms using default parameters. We ran Miniquant 1.2.2 on raw FASTQ reads of both platforms by adjusting the parameter for each platform and otherwise default parameters. Quantification for Illumina was performed using Salmon 1.10.3 using automatic libtype detection, thirty bootstrap samples, with –validateMappings, –gcBias, and –seqBias applied, a prior mean and standard deviation of the fragment length distribution of 250 and 25, respectively, and otherwise default parameters. Performance of relative quantification on the spike-in data was assessed by evaluating the Pearson correlation between estimated and true log-fold changes on the SIRVs as determined by the change in molarity between SIRV mixes E1 (E3 samples) and E2 (isoB11 samples). In addition, we performed DTE between the two SIRV mixes and calculated F1 scores of the directional predicted change of each quantification method, per platform, where we considered a prediction as negative if the transcript was called significant and had a negative log-fold change, and correspondingly for positive predictions. Performance of absolute quantification, as well as dynamic range was assessed by evaluating the Pearson correlation between true and absolute abundances for the ERCC spike-ins, evaluating the similarity between estimated logCPM values and true spike-in abundances as measured by molarity times molecular weight. Spearman’s rank correlation between Illumina GENCODE and long-read GENCODE quantifications was calculated using genes or transcripts detected (one CPM or higher in all E3 samples for long-read technologies, and the same criterion with TPM for Illumina) by at least one method in Illumina and one method in at least one of the long-read platforms, and having inferential variablity lower than 1 in the Illumina bulk dataset (when considering transcript counts). Inferential variability was computed analyzing the Salmon bootstraps with the function computeInfRV in fishpond 2.10.0 package in R. Quantification efficiency was evaluated by comparing the number of raw reads (15M) with the number of assigned counts obtained for each method. We performed DTE analysis on the transcript-level count matrices generated by each quantification method using the edgeR 4.2.2 quasi-likelihood pipeline. For each method, lowly expressed transcripts were removed using the filterByExpr function with default parameters. DTE testing was then carried out by fitting a quasi-likelihood negative binomial generalized log-linear model (glmQLFit), followed by empirical Bayes quasi-likelihood F-tests (glmQLTest). To assign confidence levels to DTE calls, we classified significant detections according to cross-method agreement. A transcript was labeled:

- High confidence if it was significantly differentially expressed (|log2(FC)|> 1 and FDR < 0.01) in one method and also significant in at least one other platform, but not detected by the same quantification method.
- Medium confidence if it was significantly differentially expressed in one method and showed a consistent direction of change (|log2(FC)|> 1) and an absolute logFC difference smaller than 2 in at least one other quantification method in the other platforms.
- Low confidence if it met the criteria for medium confidence but with an absolute logFC difference smaller than 4.

We performed DTU analysis on the transcript-level count matrices generated by each quantification method using the edgeR 4.2.2 quasi-likelihood pipeline, followed by a dedicated test for differential transcript usage at the exon level (diffSpliceDGE with test = “exon”). For each method, lowly expressed transcripts were filtered using the dmFilter function from DRIMSeq 1.32.0, with the following parameters: miṅsampṡfeaturėexpr = 3, miṅsampṡgenėexpr = 3, miṅfeaturėexpr = 10, miṅgenėexpr = 10, and miṅfeaturėprop = 0.1. After testing, we used stageR 1.30.1 to filter transcript level in order to additionally require gene level significance. Confidence levels for DTU calls were assigned following the same criteria used for DTE classification.

### Single-cell quantification

Within-platform comparisons were performed using downsampled read sets, ensuring that each quantification method for a given platform received an identical subset of reads for fair evaluation. Read depths were standardized to 60M reads,corresponding to the shallowest sample available across the three platforms. Quantification for ONT and PB was performed using Bambu, Oarfish, Kallisto, and Isosceles. Bambu c5e923dd13249ee5c7563f2e378c795024c06ee7 (Git commit hash) was run on genome alignments in the single-cell mode for both platforms with default parameters. Oarfish 0.6.3 was run on transcriptome alignments in the single-cell mode for both platforms with model coverage applied (–model-coverage), no filters and a bin width of 100. Kallisto 0.51.1 was run on raw FASTQ reads of both platforms by adjusting the parameter for each platform and otherwise default parameters. To run Kallisto in single-cell mode, we implemented a code to handle barcodes and quantify reads for each single-cell. Isosceles 0.2.1 was run on genome alignments in the single-cell mode for both platforms using default parameters. Raw Illumina single-cell reads were processed using simpleaf 0.19.0. Single-cell count data were aggregated into pseudo-bulk samples to enable comparisons following the same logic used for bulk data. Spearman’s rank correlation between Illumina GENCODE bulk profiles and long-read GENCODE pseudo-bulk profiles was computed using genes or transcripts detected (one CPM or higher in all E3 samples for long-read technologies or the same criterion for TPM for Illumina) by at least one quantification method across all long-read single-cell platforms and the Illumina bulk dataset, and having inferential variability lower than 1 in the Illumina bulk dataset (when considering transcript counts). Spearman’s rank correlation between cell-level gene counts was assessed by subsampling the same 1,000 cells from each platform. Highly variable genes were identified using the modelGeneVar function from the scran 1.32.0 R package, and the sampled genes were drawn from the top 1,000 highly variable genes for all platforms. Quantification efficiency was assessed by comparing the number of mapped and deduplicated reads with the number of counts obtained for each method. DTE and DTU tests were performed with the exact same pipeline developed for the bulk datasets. However, the classification for pseudobulk calls slightly differed. Here, a transcript was labeled:

- High confidence if it was significantly differentially expressed (|log2(FC)|> 1 and FDR < 0.01) in one method and also significant in at least one other platform including the bulk datasets, but not detected by the same quantification method.
- Medium confidence if it was significantly differentially expressed in one method and showed a consistent direction of change (|log2(FC)|> 1) and an absolute logFC difference smaller than 2 in at least two other quantification methods in other platforms, including the bulk datasets.
- Low confidence if it met the criteria for medium confidence but with an absolute logFC difference smaller than 4.

## Case study

In our case study, we calculated the Wasserstein distance for each gene that exhibited non-zero counts in all technologies for each platform. For every gene, transcript-level proportions were derived by normalizing each transcript’s counts by the total counts across all transcripts of that gene. Using these transcript proportions, we computed the Wasserstein distance with a uniform cost matrix, implemented via the Wasserstein function from the transport 0.15-4 R package.

## Data availability

Raw data of all single-cell and bulk samples are openly available from ArrayExpress (single-cell accession: E-MTAB-16805; bulk accession: E-MTAB-16791). Processed data, including full-depth quantifications of our recommended methods for all samples and downsampled quantifications of all methods for all samples, are available from Zenodo under an MIT license (Botta et al. 2026a).

## Code availability

A Snakemake pipeline reproducing all of our analyses, including figures, is available under an MIT license from Github (Botta et al. 2026b). All code has additionally been archived under an MIT license in Zenodo (Botta et al. 2026a).

## Reproducibility

All analyses were run via containerized conda environments managed by a Snake-make workflow using apptainer 1.4.5 and snakemake 9.11.2. All computations were performed on Ubuntu 22.04 with AMD EPYC 7763 CPUs. For timing and memory comparisons, all single-cell methods were given access to 24 cores, and all bulk methods were given access to six cores.

## Declarations

G.B., S.H., T.C., K.R., S.P., J. R. C., K. S., Z. M. N. C. D. N., S. J. C., and R. S. are, or were at the time their contribution took place, employees and/or shareholders of the F. Hoffmann-La Roche AG. A. B., Z. M., and W. S. are, or were at the time their contribution took place, employees and shareholders of Genentech, Inc. All remaining authors declare no competing interests.

