## Supplementary information for "Benchmarking long-read RNA-seq across modalities, methods, and sequencing depth in iNeurons"

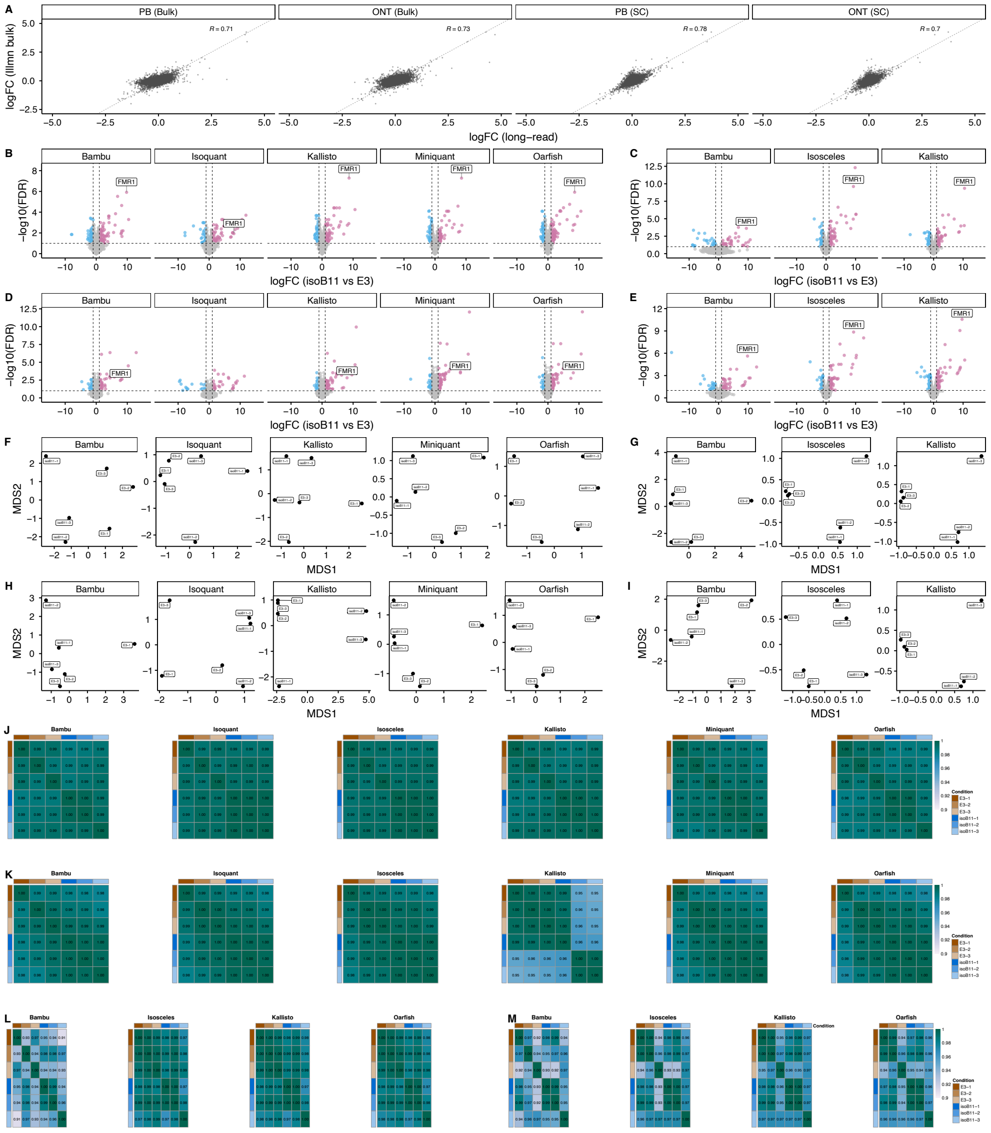


**Fig. S1**: **The FXS system demonstrates a clear separation between the two conditions. A.** Estimated edgeR gene-level log-fold change concordances between Illumina and all long-reach technologies, filtered for genes that have one or more CPM/TPM in all shown quantifications. **B-E.** DGE analysis between E3 and IsoB11 samples with all the remaining methods not shown in Fig. 1C for all the datasets (B: Oxford Nanopore Technologies (ONT) bulk, C: ONT pseudobulk, D: Pacific Biosciences (PB) bulk, E: PB pseudobulk, respectively). **F-I.** Multidimensional scaling (MDS) plots for all quantification methods not shown in Fig. 1D for transcript counts in all datasets (F: ONT bulk, G: ONT pseudobulk, H: PB bulk, I: PB pseudobulk, resepectively). **J-M.** Pearson correlation analysis across all samples for transcript counts in all datasets (J: ONT bulk, K: ONT pseudobulk, L: PB bulk, M: PB pseudobulk, respectively).

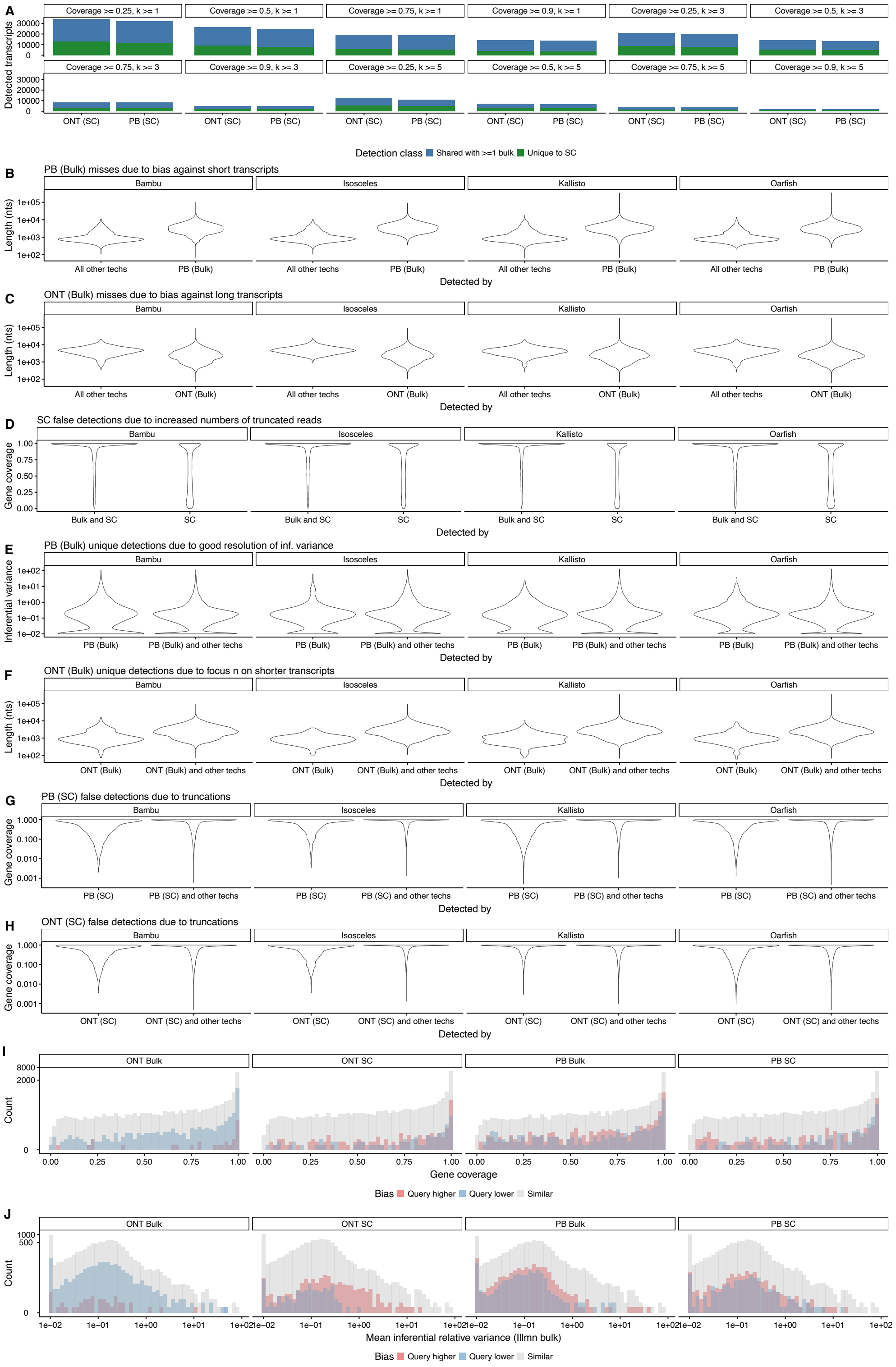

A
Detected transcripts
Coverage >= 0.25, k >= 1
Coverage >= 0.5, k >= 1
Coverage >= 0.75, k >= 1
Coverage >= 0.9, k >= 1
Coverage >= 0.25, k >= 3
Coverage >= 0.5, k >= 3
Coverage >= 0.75, k >= 3
Coverage >= 0.9, k >= 3
Coverage >= 0.25, k >= 5
Coverage >= 0.5, k >= 5
Coverage >= 0.75, k >= 5
Coverage >= 0.9, k >= 5
ONT (SC)
PB (SC)
Detection class
Shared with >=1 bulk
Unique to SC
B
PB (Bulk) misses due to bias against short transcripts
Bambu
Isosceles
Kallisto
Oarfish
Length (nts)
All other techs
PB (Bulk)
Detected by
C
ONT (Bulk) misses due to bias against long transcripts
All other techs
ONT (Bulk)
D
SC false detections due to increased numbers of truncated reads
Gene coverage
Bulk and SC
SC
E
PB (Bulk) unique detections due to good resolution of inf. variance
Inferential variance
PB (Bulk)
PB (Bulk) and other techs
F
ONT (Bulk) unique detections due to focus n on shorter transcripts
ONT (Bulk)
ONT (Bulk) and other techs
G
PB (SC) false detections due to truncations
PB (SC)
PB (SC) and other techs
H
ONT (SC) false detections due to truncations
ONT (SC)
ONT (SC) and other techs
I
ONT Bulk
ONT SC
PB Bulk
PB SC
Count
Gene coverage
Bias
Query higher
Query lower
Similar
J
Mean inferential relative variance (Illmn bulk)

**Fig. S2**: **Platform or technology-specific biases are consistent between quantification methods, and, conditional on detection, length is the main factor driving further quantification differences between platforms or technologies. We only show methods throughout that are available for both single-cell and bulk data. A.** Number of detected transcripts in single-cell technologies that are shared with at least one bulk technology, or unique to single-cell, stratified by filtering options which only retain transcripts that have a minimum gene coverage and are part of a gene that has a minimum of $k$ other transcripts detected. **B.** Length differences between transcripts detected by all other methods and transcripts detected by PB (Bulk) (shown for all methods that are available for bulk and single-cell, since Figure 2 uses different methods for bulk and single-cell). **C.** Length differences between transcripts detected by all other methods and transcripts detected by ONT (Bulk) (shown for all methods that are available for bulk and single-cell, since Figure 2 uses different methods for bulk and single-cell). **D.** Gene coverage (see **Methods**) differences between transcripts detected by both single-cell technologies, but none of the bulk technologies (shown for all methods that are available for bulk and single-cell, since Figure 2 uses different methods for bulk and single-cell). **E.** Mean inferential relative variance in Illumina (Bulk) (see **Methods**) differences between transcripts detected by PB (Bulk) uniquely compared to transcripts shared with at least one other technology or platform. (shown for all methods that are available for bulk and single-cell, since Figure 2 uses different methods for bulk and single-cell). **F.** Length differences between transcripts detected by ONT (Bulk) uniquely compared to transcripts shared with at least one other technology or platform. (shown for all methods that are available for bulk and single-cell, since Figure 2 uses different methods for bulk and single-cell). **G.** Gene coverage (see **Methods**) differences between transcripts detected by ONT (SC) uniquely compared to transcripts shared with at least one other technology or platform. (shown for all methods that are available for bulk and single-cell, since Figure 2 uses different methods for bulk and single-cell). **H.** Gene coverage (see **Methods**) differences between transcripts detected by PB (SC) uniquely compared to transcripts shared with at least one other technology or platform. (shown for all methods that are available for bulk and single-cell, since Figure 2 uses different methods for bulk and single-cell). **I.** Histogram of gene coverage of transcripts that are similar, or notably lower or higher in estimated abundance (see **Methods**) between a query technology and all other technologies, stratified by query technology. **J.** Histogram of mean inferential relative variance in Illumina (Bulk) (see **Methods**) of transcripts that are similar, or notably lower or higher in estimated abundance (see **Methods**) between a query technology and all other technologies, stratified by query technology.

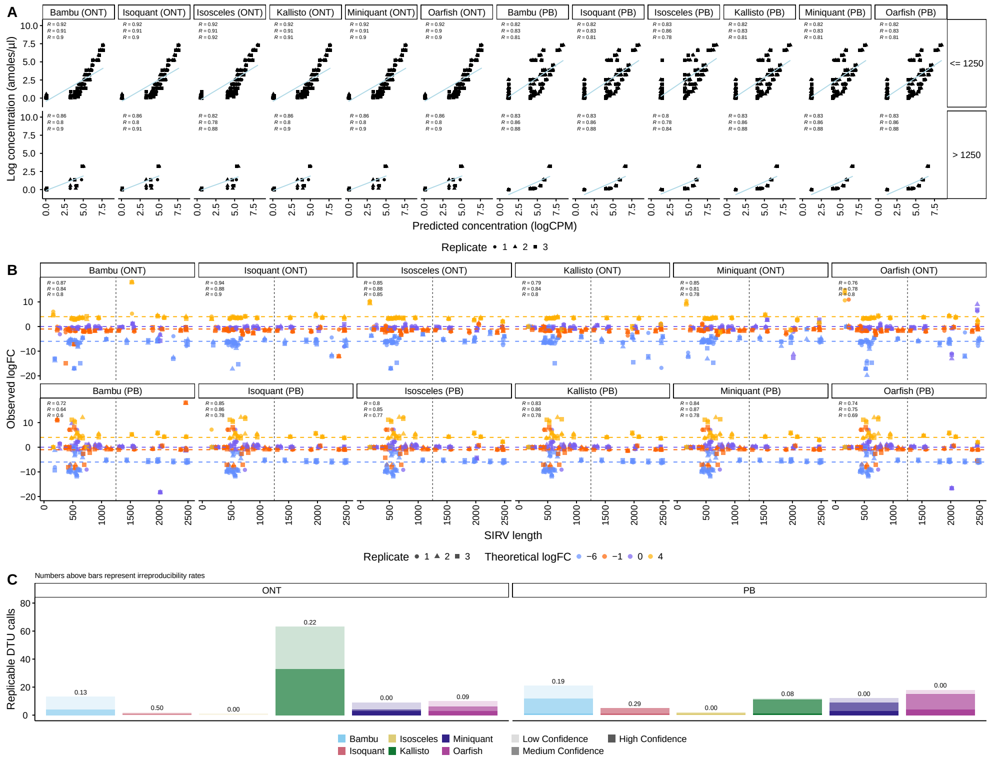


**Fig. S3**: **Bulk DTU performance, overlap between DTE and DTU calls between methods, and Illumina-lrRNA-seq correlations mirror spike-in and DTE performance results. A.** Observed logCounts Per Million (CPM) (y-axis) versus expected amoles/$\mu$ (x-axis) for ERCC spike-in molecules, stratified by transcript length, across different quantification methods and technologies. Numbers in the top-left of each subplot indicate Spearman's rank correlation coefficients calculated across all SIRVs in each E3 replicate. Overall, ONT exhibits higher quantification accuracy than PB, particularly for transcripts shorter than 1.25 kb, where PB shows substantial underdetection. **B.** Observed (y-axis) versus expected (x-axis) logFC of SIRVs, stratified by length, for each quantification method and sequencing platform. Numbers in the top-left of each subplot indicate Spearman's rank correlation coefficients calculated across all SIRVs in each E3 replicate. PB exhibits misquantification for SIRVs shorter than 1.25kb. Among methods, Isoquant achieves the highest correlation with expected logFC values. **C.** Replicable DTU calls for each quantification method across sequencing platforms at FDR = 0.01. Confidence is assessed through cross-method and cross-platform agreement (see **Methods**).

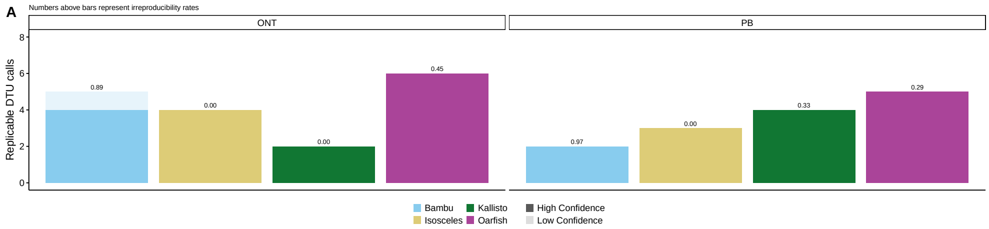


**Fig. S4**: **A.** Replicable DTU calls for each quantification method across sequencing platforms at FDR = 0.01. Confidence is assessed through cross-method and cross-platform agreement (see **Methods**).

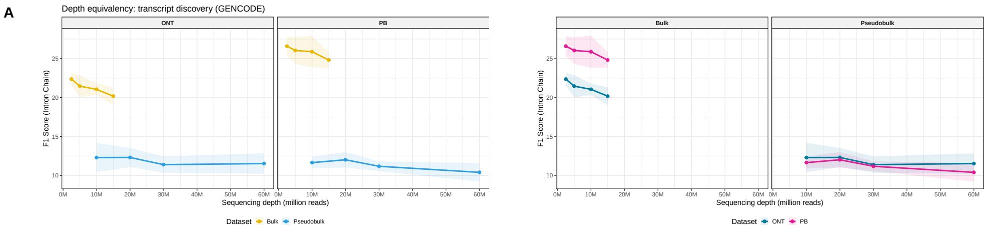


**Fig. S5**: **Establishing depth equivalencies between both technologies and platforms was not possible on GENCODE transcript re-discovery due to strong outperformance by bulk platforms compared to single-cell (far left, middle left), outperformance of PB bulk compared to ONT bulk (middle right), and broadly equivalent performance between the two single-cell technologies (far right). A.** Depth equivalency calculated on the F1 score of attempting to re-discover selected GENCODE transcripts expressed across technologies and platforms (see **Methods**). Far left: ONT bulk compared to ONT pseudobulk. Middle left: PB bulk compared to PB pseudobulk. Middle right: ONT bulk compared to PB bulk. Far right: ONT pseudobulk compared to PB pseudobulk.
